# Endometrial cancer progression driven by PTEN-deficiency requires miR-424(322)^∼^503

**DOI:** 10.1101/2025.04.07.647575

**Authors:** Maria Vidal-Sabanés, Núria Bonifaci, Raúl Navaridas, Joaquim Egea, Mario Encinas, Ruth Rodriguez-Barrueco, Jose M. Silva, Xavier Matias-Guiu, David Llobet-Navas, Xavier Dolcet

**Affiliations:** Developmental and Oncogenic Signaling Group. Departament de Ciències Mèdiques Bàsqieus/Medicina Experimental. Universitat de Lleida. Institut de Recerca Biomèdica de Lleida, IRBLleida. Lleida, Spain; Molecular Mechanisms and Experimental Therapy in Oncology-Oncobell Program. Bellvitge Biomedical Research Institute (IDIBELL), L’Hospitalet de Llobregat, Barcelona, Spain; Centro de Investigación Biomédica en Red de Cáncer (CIBERONC), Instituto de Salud Carlos III, (ISCIII) - Madrid, Spain; Herbert Irving Comprehensive Cancer Center; Vagelos College of Physicians and Surgeons, Columbia University Irving Medical Center, New York, NY, 10032, USA; Department of Pathology and Experimental Therapy, School of Medicine, Universitat de Barcelona. L’Hospitalet de Llobregat, Spain; Department of Pathology, Icahn School of Medicine at Mount Sinai, New York, NY, 10029, USA; Oncologic Pathology Group. Departament de Ciències Mèdiques Bàsiques, Universitat de Lleida. Institut de Recerca Biomèdica de Lleida, IRBLleida. Lleida, Spain

**Keywords:** miR-424(322)^∼^503, miR-424, miR-503, PI3K/AKT, PTEN, TGFβ, endometrial cancer

## Abstract

Endometrial cancer is the most frequent type of cancer in the female reproductive tract. Loss-of-function alterations in PTEN, leading to enhanced PI3K/AKT activation, are among the most frequent molecular alterations in endometrial cancer. Increased PI3K/AKT signaling resulting from PTEN loss promotes cellular proliferation and confers resistance to TGFβ-mediated apoptosis, a key regulator of endometrial homeostasis. In this study, we have analyzed the role of miRNAs in driving these altered cellular responses. A comprehensive transcriptomic analysis of miRNA expression revealed the upregulation of several miRNAs caused by PTEN deficiency and/or TGFβ stimulation. The miR-424(322)^∼^503 cluster drew our attention due to its involvement in regulating apoptosis and proliferation. However, miR-424(322)^∼^503 cluster has a paradoxical role in cancer, exhibiting either oncogenic and tumor suppressive functions depending on cell type or context. To ascertain the function of miR-424(322)^∼^503 in endometrial carcinogenesis caused by PTEN deficiency, we generated a double Pten/miR-424(322)^∼^503 knock-out mice. We demonstrate that loss of miR-424(322)^∼^503 impairs proliferation of both wild type or *Pten* deficient endometrial organoids by interfering with growth factor and PI3K/AKT signaling. Furthermore, the absence of miR-424(322)^∼^503 restores TGFβ-induced apoptosis, which is otherwise compromised by PTEN deficiency. *In vivo*, *Pten*/miR-424(322)^∼^503 knock-out mice exhibit reduced endometrial cancer progression compared to *Pten* deficient mice through a cell-autonomous mechanism.

## INTRODUCTION

Endometrial cancer (EC) is the most common malignancy affecting the female reproductive tract ^1^ ^2^, with its incidence steadily rising worldwide. The molecular mechanisms underlying EC development are complex and multifaceted, involving key signaling pathways such as PI3K/AKT and TGFβ/Smad, which play crucial roles in maintaining endometrial homeostasis. Dysregulation of these pathways significantly contribute to the initiation and progression of EC. Among these alterations, loss-of-function mutations in *PTEN* are the most frequent molecular events in EC. These mutations lead to hyperactivation of the PI3K/AKT signaling pathway, promoting uncontrolled cellular proliferation and survival. The pivotal role of *PTEN* mutations in EC development has been extensively validated through genetically engineered mouse models. These include models with heterozygous *Pten* deletions, which mimic early-stage EC ^3-5^, *in vivo* CRISPR-Cas9 gene editing ^6^ as well as conditional *Pten* deletions in the endometrium, which provide a more tissue-specific understanding of its role in tumorigenesis ^7^.

In parallel, the disruption of TGFβ signaling has emerged as another critical factor in endometrial carcinogenesis ^8^ ^9^. TGFβ signaling, mediated through the Smad pathway, is essential for maintaining growth inhibition and tissue integrity in the endometrium. However, EC frequently exhibits impaired TGFβ signaling, which undermines its tumor-suppressive effects. This impairment not only results in the loss of growth inhibition but also facilitates the acquisition of invasive and metastatic phenotypes, correlating with poor clinical outcomes ^10-13^. The significance of TGFβ/Smad signaling in EC has been further elucidated through advanced genetic models. Conditional abrogation of TGFβ signaling components, such as the deletion of *T*β*RI* (the TGFβ receptor I) ^14^, has demonstrated its essential role in preventing tumorigenesis. Moreover, models with the combined deletion of *SMAD2* and *SMAD3 ^15^*, or dual deletion of *T*β*RI* alongside *PTEN* in the endometrium, highlight the synergistic impact of these pathways in driving endometrial cancer progression ^16^. These findings underscore the intricate interplay between the PI3K/AKT and TGFβ/Smad pathways in endometrial carcinogenesis, offering valuable insights into potential therapeutic strategies targeting these molecular alterations.

The miR-424(322)^∼^503 cluster encodes two broadly conserved miRNAs, miR-424 (322 in mouse) and miR-503, which belong to the miR-15/107 family ^17^ ^18^. These miRNAs play a critical role in regulating various cancer-related cellular processes, including proliferation, differentiation, cellular plasticity, and apoptosis ^19^. The miR-424(322)^∼^503 cluster has been reported to exhibit altered expression in multiple cancer types, being either upregulated or downregulated depending on the context. Functionally, the role of the miR-424(322)^∼^503 cluster in cancer is paradoxical, as it can act as either an oncogene or a tumor suppressor, depending on the cell type and microenvironment ^20-22^. This duality highlights the complexity of its regulatory mechanisms and underscores its context-dependent functions in tumorigenesis. The expression of miR-424(322)^∼^503 is tightly regulated by various signaling pathways and transcription factors. Notably, the TGFβ signaling pathway establishes a dynamic feedback regulatory loop with miR-424(322)^∼^503. On one hand, TGFβ signaling transcriptionally induces the expression of the miR-424(322)^∼^503 cluster ^23^ ^24^. On the other hand, miR-424(322)^∼^503 directly targets key components of the TGFβ/Smad signaling cascade, including SMURF2 and SMAD7 ^25^, which act as negative regulators of TGFβ signaling. Additionally, miR-424(322)^∼^503 can modulate the pathway by targeting core signaling mediators such as SMAD2 ^26^ and SMAD3 ^27^. Through these interactions, miR-424(322)^∼^503 exerts significant control over the activation and downstream effects of TGFβ signaling.

The role of the miR-424(322)^∼^503 cluster in endometrial cancer (EC) has been minimally explored, with most studies focusing on its function in human cancer cell lines. However, no investigations to date have addressed the *in vivo* function of the miR-424(322)^∼^503 cluster in EC using genetically modified mouse models. Similar to other malignancies, the role of miR-424(322)^∼^503 in EC remains controversial i.e., while the majority of studies suggest a tumor-suppressive function for miR-424(322)^∼^503 in EC ^28-34^, others have recently provided evidence supporting its oncogenic role ^35^. Building on these findings, we aimed to investigate the role of the miR-424(322)^∼^503 cluster in PTEN-loss-driven EC. By modeling complex experimental genetic combinations our study demonstrates that deletion of miR-424(322)^∼^503 significantly reduces cell proliferation in both normal and PTEN-deficient endometrial organoids by interfering with insulin signaling and the PI3K/AKT pathway. Additionally, miR-424(322)^∼^503 deficiency restores sensitivity to TGFβ-induced apoptosis in *Pten*-deficient organoids. Finally, using a double Pten/miR-424(322)^∼^503 knockout mouse model, we provide compelling *in vivo* evidence that the absence of miR-424(322)^∼^503 dramatically reduces endometrial neoplasia driven by PTEN deficiency. This reduction occurs through a cell-autonomous mechanism, highlighting the miR-424(322)^∼^503 cluster as a critical regulator of endometrial carcinogenesis in the context of PTEN loss.

## RESULTS

### Genome-wide miRNA profiling identifies miR-322 as a potential key mediator of PTEN loss signaling

In recent years, our laboratory has focused on unraveling the mechanisms of endometrial carcinogenesis through the use of both *in vivo* and *in vitro* models. Among these, we have employed an inducible conditional *Pten* knock-out murine model (Cre:ER^(T)+/-^Pten^F/F^), which reliably develops endometrial carcinomas ^36^, and three-dimensional (3D) cultures of endometrial organoids ^37^. Importantly, using endometrial organoids derived from the conditional *Pten* knockout model, we demonstrated that *Pten* deficiency induces resistance to TGFβ-induced cell death ^38^. This finding underscores the pivotal role of PTEN in modulating the cellular response to TGFβ signaling, a pathway critical for maintaining endometrial homeostasis. To further elucidate the mechanisms by which PTEN loss counteracts TGFβ-induced cell death and to examine the potential involvement of microRNAs in this process, we performed small RNA sequencing (miRNA-seq) on endometrial organoids derived from both normal epithelial cells (Cre:ER^(T)-/-^Pten^F/F^; Wt) and *Pten*-deficient cells (Cre:ER^(T)+/-^Pten^F/F^) treated with tamoxifen **(Figure 1A)**. Interestingly, the analysis of the genetic miRNA program associated to *Pten* loss revealed a significant enrichment of several miRNAs in *Pten*-deficient samples compared to Wt controls (56 differentially expressed miRNAs, log_2_ FC > abs(1.5) plus adj. FDR p<0.05), highlighting the potential involvement of miRNAs in the cellular changes associated with *Pten* loss. Among the upregulated miRNAs, we identified the 10 with the most statistically significant p-values **(Figure 1B, Supplementary Table S1)**. Notably, a steady increase in miR-322-5p (hereafter referred to as miR-322 in mice or miR-424 in humans) was observed in *Pten*-deficient organoids. Interestingly, miR-424 has previously been linked to the regulation of the PI3K/AKT pathway and TGFβ signaling ^23-27^, highlighting its potential significance in the underlying biological processes associated to PTEN loss. Importantly, miR-424 is genomically located on the Xq26.3 region in the X chromosome in close proximity to miR-503 in both mice and humans. The two miRNAs are expressed as a cluster ^39^, and their co-regulation suggests a coordinated role in cellular function. Moreover, the Xq26.3 region, which includes the lncRNA H19X (MIR503HG) and the syntenic cluster of miR-424 and miR-503, has been consistently reported as altered in various cancers, underscoring its significance in tumorigenesis. Both miR-424 and miR-503 belong to the miR-15/107 family of microRNAs, which is highly conserved across mammals and plays a pivotal role in regulating essential cellular processes such as proliferation, apoptosis, and differentiation ^18^ ^19^. Collectively, this evidence suggests that the miR-424(322)^∼^503 cluster may play a critical role in endometrial cancer progression driven by PTEN loss. Hence, to validate our miRNA_seq findings, we performed RT-qPCR to measure the expression levels of miR-322 and miR-503 in Wt and *Pten* KO endometrial organoids. Consistent with the sequencing data, both miR-322 and miR-503 showed a marked increase in expression in *Pten*-deficient organoids **(Figure 1C)**, further supporting their involvement in the molecular mechanisms associated with PTEN loss. Remarkably, a direct comparison of the expression levels of miR-322 and miR-503 within this cluster revealed an unbalanced expression ratio, with miR-322 being expressed at significantly higher levels than miR-503 **(Figure 1D-E)**. This observation was further corroborated by an analysis of relative miR-424 and miR-503 expression values derived from RNA-seq data in the TCGA-UCEC database ^40^. The analysis demonstrated a strong positive correlation between the expression of these paralogs **(Figure 1F)**, accompanied by a stability imbalance characterized by markedly higher expression of miR-424 compared to miR-503 **(Figure 1G)**. This observation aligns with previous studies reporting a higher turnover rate of miR-503, attributed to the nucleotide composition within its seed region and the 3’ miRNA end ^41^. Such molecular instability appears to be a shared feature among other members of the miR-15/107 family and is thought to play a critical role in fine-tuning their expression levels in specific cellular contexts. Prompted by this data, and given the established role of the miR-424(322)^∼^503 cluster in regulating cancer-related cellular processes, we decided to focus our investigation on the function of this cluster of miRNAs in *Pten*-loss-induced carcinogenesis.

**Figure 1.**
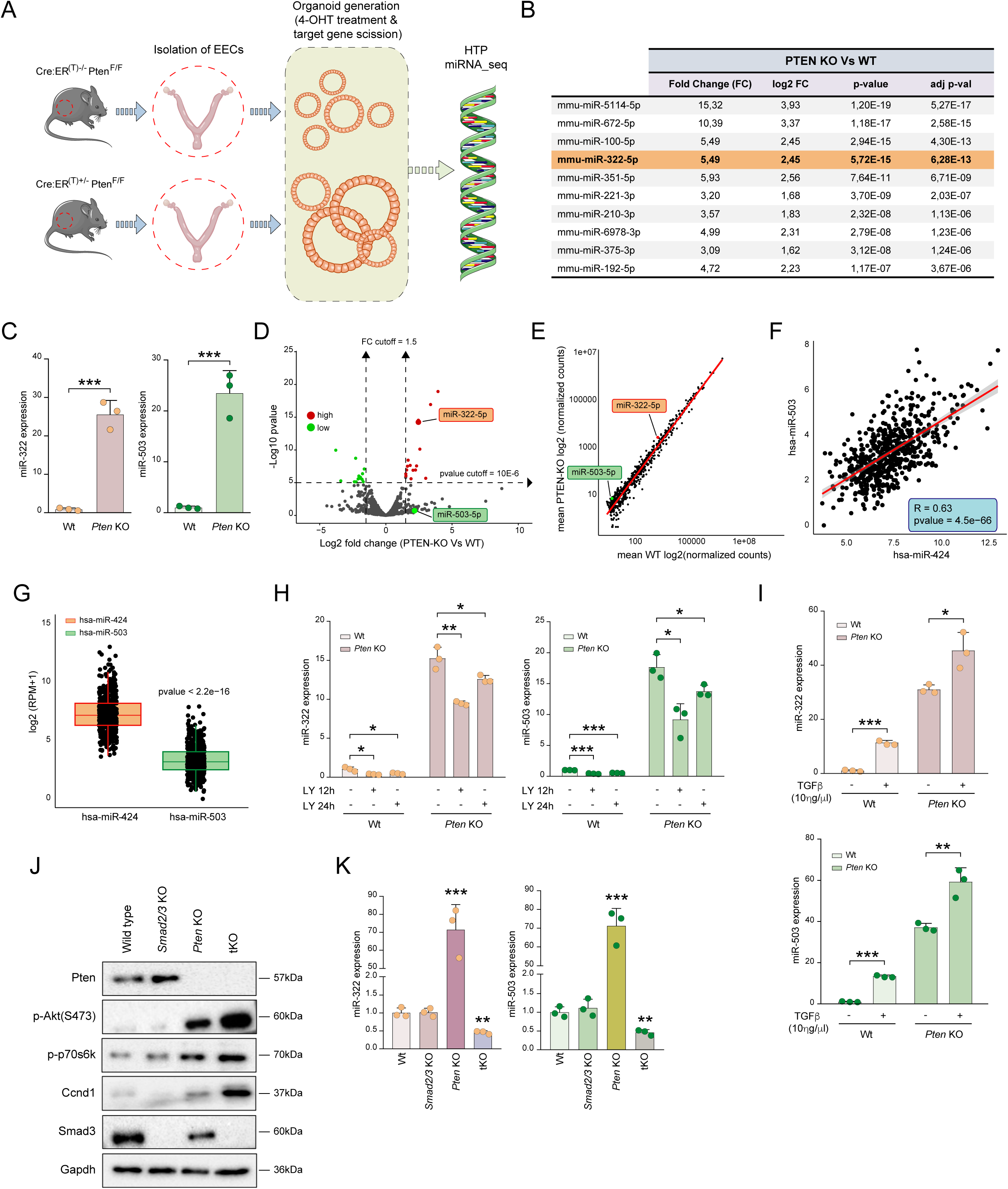
Loss of PTEN increases miR-322^∼^503 expression via TGFβ and PI3K/AKT pathways. **(A)** Schematic illustration of the experimental design. **(B)** Top 10 miRNAs with significantly increased expression in Cre:ER^(T)+/-^Pten^F/F^ organoids (*Pten* KO) organoids compared to Cre:ER^(T)-/-^Pten^F/F^ (Wt) organoids. Data include expression values from miRNA-seq, p-values, and adjusted p-values. **(C)** Relative expression of miR-322 and miR-503 in Wt and *Pten* KO mouse endometrial organoids. **(D)** Volcano plot showing the distribution of miRNAs according to p-values and fold changes between *Pten* KO and Wt conditions. **(E)** Scatterplot showing the correlation of miRNA-seq data, with average expression values from *Pten* KO samples (Y-axis) plotted against Wt samples (X-axis). **(F)** Pearson correlation analysis of miR-424 and miR-503 expression using data from the TCGA-UCEC database (n=575). **(G)** Box plot comparing the relative expression of miR-503 and miR-424 in the TCGA-UCEC database. **(H)** Relative expression of miR-322 and miR-503 in Wt and *Pten* KO 3D mouse organoids treated with 20μM LY-294002 for 12 and 24 hours. **(I)** Relative expression of miR-322 and miR-503 in Wt and *Pten* KO mouse endometrial organoids after treatment with 10ηg/μL TGFβ for 16 hours. **(J)** Representative western blot images of signaling proteins: p-Pten, p-Akt^Ser473^, p-p70s6k, Ccnd1 and Smad3. The analysis was performed on Cre:ER^(T)-/-^Pten^F/F^Smad2/3^F/F^ (Wt), Cre:ER^(T)+/-^Pten^F/F^Smad2/3^+/+^ (*Pten* KO), Cre:ER^(T)+/-^ Pten^+/+^Smad2/3^F/F^ (*Smad2/3* KO), and Cre:ER^(T)+/-^Pten^F/F^Smad2/3^F/F^ (*Pten* and *Smad2/3* triple knockout, i.e., tKO). Membranes were re-probed with Gapdh antibody as a protein loading control. **(K)** Relative expression of miR-322 and miR-503 in Wt, *Pten* KO, *Smad2/3* KO, and tKO mouse endometrial organoid cultures. Data are presented as mean ±SD, by t-test analysis, *p<0.05; **p<0.01; ***p<0.001.

### Pharmacologic and genetic engineering studies unveil PI3K/AKT and TGF**b** signaling as regulators of miR-322^∼^503 expression

Our miRNA-seq analysis of endometrial organoids derived from Cre:ER^(T)-/-^Pten^F/F^ (Wt) and Cre:ER^(T)+/-^Pten^F/F^ (*Pten* KO) mice revealed that PTEN deficiency significantly upregulated the expression of the miR-322^∼^503 cluster. Given the well-established role of the PI3K/AKT signaling pathway in mediating cellular responses to *Pten* loss, we next investigated whether the observed upregulation of miR-322^∼^503 was driven by enhanced PI3K/AKT activity. To this end, Wt and *Pten* KO organoids were treated with the PI3K inhibitor LY-294002, a synthetic chemical compound that functions as a potent phosphoinositide 3-kinase (PI3K) inhibitor, for 12 and 24 hours, followed by measurement of miR-322 and miR-503 expression. Notably, LY-294002 treatment led to a significant reduction in the expression of both miRNAs in Wt and *Pten* KO organoids **(Figure 1H)**, confirming that PI3K/AKT signaling contributes to the regulation of miR-322^∼^503 in this context.

Previous studies from our laboratory have demonstrated that the expression of the miR-424(322)^∼^503 cluster is induced by activation of the TGFβ/Smad signaling pathway in the mammary epithelium ^23^ ^24^. Additionally, we have shown that *Pten* loss leads to constitutive nuclear translocation of Smad2/3 in endometrial organoids ^42^, potentially indicating that TGFβ signaling may be involved in miR-424(322)^∼^503 expression. Notably, treatment of wild-type organoids with TGFβ led to a significant upregulation of these miRNAs, further supporting this hypothesis **(Figure 1I)**. These findings provided a strong foundation to investigate whether the observed increase in miR-322 and miR-503 expression in *Pten*-deficient cells is mediated by Smad2/3 activity. To address this, we analyzed miR-322 and miR-503 expression across a panel of genetically engineered mouse-derived endometrial organoids with specific genetic combinations of *Pten* and *Smad2/3* deletions. The genotypes included Cre:ER^(T)-/-^Pten^F/F^Smad2/3^F/F^ (Wt), Cre:ER^(T)+/-^ Pten^F/F^Smad2/3^+/+^ (*Pten* KO), Cre:ER^(T)+/-^Pten^+/+^Smad2/3^F/F^ (*Smad2/3* KO), and Cre:ER^(T)+/-^ Pten^F/F^Smad2/3^F/F^ (*Pten* and *Smad2/3* triple knockout, i.e., tKO) **(Figure 1J)**. Quantitative analysis of miR-322 and miR-503 expression revealed that the absence of S*mad2/3* in *Pten*-deficient organoids completely abolished miR-322 and miR-503 expression **(Figure 1K)**. These results strongly suggest that the transcriptional activity of SMAD2/3, driven by its nuclear translocation in the context of PTEN loss, is indispensable for the upregulation of miR-322 and miR-503. These findings highlight a dual regulatory mechanism in which PTEN loss and TGFβ signaling converge to modulate the expression of the miR-424(322)^∼^503 cluster, with PI3K/AKT signaling playing a central role in this process.

### miR-322∼503 deficiency impairs proliferation in *Pten*-deficient and Wt endometrial organoids

PTEN loss induces hyperactivation of PI3K/AKT signaling leading to an aberrant glandular proliferation and endometrial cancer (EC) initiation. To investigate the role of the miR-424(322)^∼^503 cluster in PTEN-deficient EC, we generated a double knockout model by crossing miR-322^∼^503 knockout mice (miR KO) ^21^ with a *Pten*-conditional knockout strain (*Pten* KO) ^33^. Epithelial endometrial cells were isolated from Cre:ER(^T)-/-^Pten^F/F^miR-322^∼^503^+/+^ (Wt), Cre:ER^(T)-/-^ Pten^F/F^miR-322^∼^503^-/-^ (miR KO), Cre:ER^(T)+/-^Pten^F/F^miR-322^∼^503^+/+^ (*Pten* KO), and Cre:ER^(T)+/-^ Pten^F/F^miR-322^∼^503^-/-^ (double knockout, i.e., dKO). and cultured in a three-dimensional organoid system to assess the functional impact of miR-322^∼^503 deficiency on glandular proliferation. Organoid cultures revealed that the absence of miR-322^∼^503 completely abrogated the increased organoid size associated with *Pten* loss **(Figure 2A)**. Quantitative analysis demonstrated a significant reduction in glandular perimeter in dKO organoids compared to *Pten* KO organoids, as well as in miR KO organoids compared to Wt organoids **(Figure 2B)**. These findings suggest that miR-322^∼^503 is required for the proliferation of both *Pten*-deficient and Wt endometrial epithelial cells.

**Figure 2.**
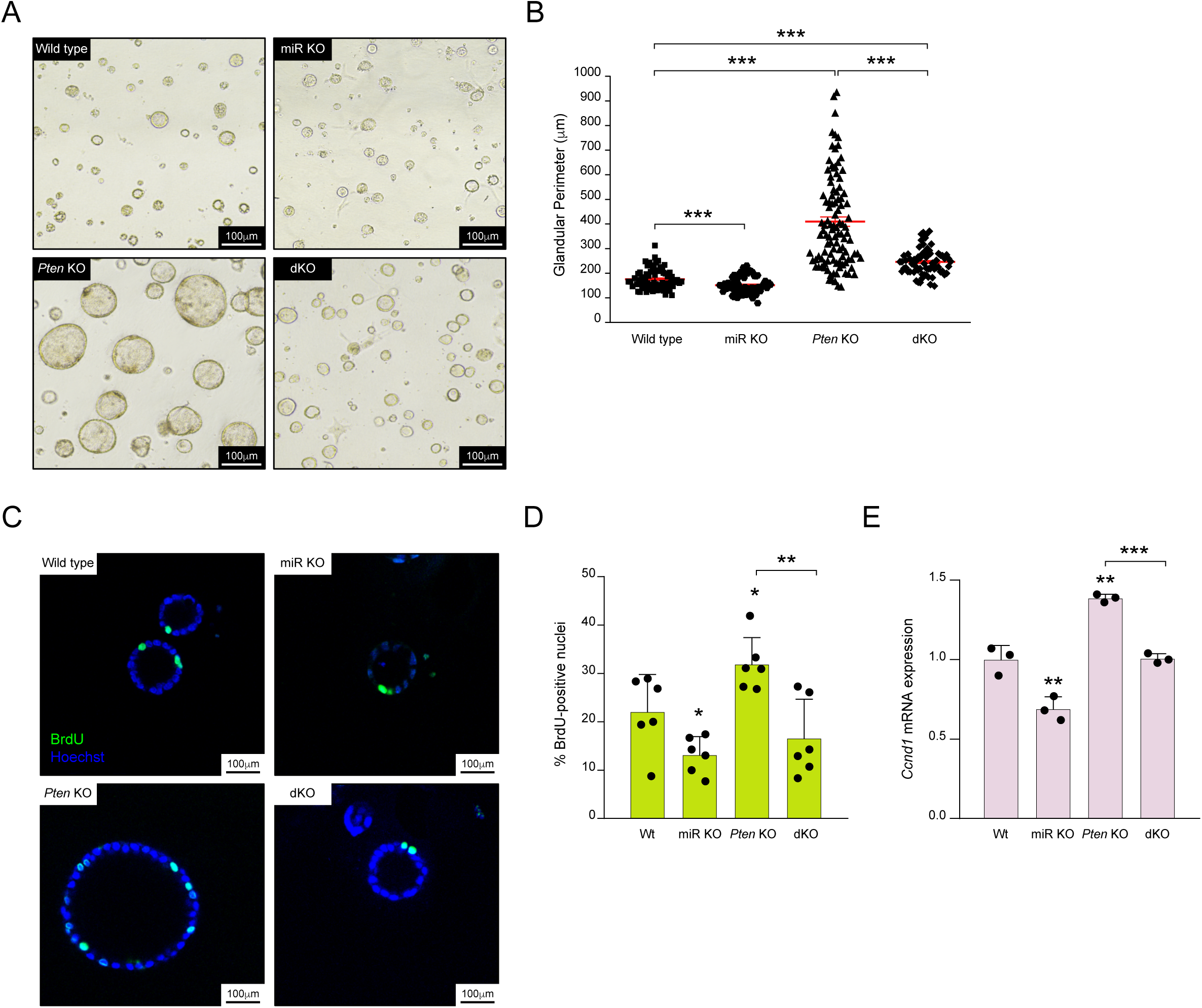
MiR-322^∼^503 is essential for the proliferation of wild type and *Pten*-deficient endometrial cells. **(A)** Representative images of endometrial organoids from Cre:ER^(T)-/-^ Pten^F/F^miR-322^∼^503^+/+^ (Wt), Cre:ER^(T)-/-^Pten^F/F^miR-322^∼^503^-/-^ (miR KO), Cre:ER^(T)+/-^Pten^F/F^miR-322^∼^503^+/+^ (*Pten* KO), and Cre:ER^(T)+/-^Pten^F/F^miR-322^∼^503^-/-^ (dKO) mice. **(B)** Quantification of glandular perimeter for the organoids. Scale bar: 100μm (20X magnification). **(C)** Representative confocal images showing 5-Bromo-2’-deoxyuridine (BrdU)-positive nuclei in Wt, miR KO, *Pten* KO, and dKO mouse endometrial organoids. **(D)** Quantification of BrdU-positive nuclei as a proportion of total nuclei, stained with Hoechst. Scale bar: 20μm (60X magnification). **(E)** Relative expression of *Ccnd1* mRNA in Wt, miR KO, *Pten* KO, and dKO mouse endometrial organoids. Data are presented as mean ±SD, by t-test analysis, *p<0.05; **p<0.01; ***p<0.001.

To further evaluate the impact of miR-322^∼^503 deficiency on cellular proliferation, we performed a 5-bromo-2’-deoxyuridine (BrdU) incorporation assay. BrdU-positive cells were significantly reduced in miR KO organoids compared to Wt organoids, and in dKO organoids compared to *Pten* KO organoids, indicating a decreased proliferation rate in the absence of miR-322^∼^503 **(Figure 2C-D)**. To corroborate these findings, we studied the expression levels of Cyclin D1 (*Ccnd1*), a well-known proto-oncogene in EC which expression we have previously demonstrated to be enhanced upon PTEN deletion in the mouse endometrium ^43^. Cyclin D1 expression, a marker of cell cycle progression, was analyzed by RT-qPCR and we found that Cyclin D1 levels were significantly lower in dKO organoids compared to *Pten* KO organoids, and in miR KO organoids compared to Wt organoids **(Figure 2E)**. These results confirm that miR-322^∼^503 deficiency leads to a reduction in cell cycle activity, thereby impairing proliferation in both *Pten*-deficient and Wt endometrial epithelial organoids. Altogether, these data highlight the essential role of the miR-322^∼^503 cluster in driving cellular proliferation in endometrial organoids, both in the context of PTEN loss and under normal conditions. This underscores the potential of targeting miR-322^∼^503 as a therapeutic strategy in PTEN-driven endometrial carcinogenesis.

### miR-322^∼^503 deficiency negatively regulates gene expression of genes involved in response to growth factors

Our findings collectively indicate that miR-322^∼^503 functions as a pivotal regulator of PI3K/AKT-driven epithelial cell expansion in the endometrium. This highlights its critical role in promoting cellular proliferation under both normal and PTEN-deficient conditions. The biological functions of miRNAs primarily arise from their capacity to fine-tune complex transcriptional networks by altering the stability of mRNAs ^44^. Hence, to further elucidate the genome-wide transcriptional changes associated with miR-322^∼^503 targeted deletion and to provide a robust framework to investigate the landscape of downstream genetic networks and pathways modulated by miR-322^∼^503, we conducted deep bulk RNA sequencing. This approach allowed us to profile the genetic programs influenced by miR-322^∼^503 loss across various endometrium-derived organoid genotypes. Specifically, we analyzed organoids derived from Cre:ER^(T)-/-^Pten^F/F^miR-322^∼^503^+/+^ (Wt), Cre:ER^(T)-/-^Pten^F/F^miR-322^∼^503^-/-^ (miR KO), Cre:ER^(T)+/-^Pten^F/F^miR-322^∼^503^+/+^ (*Pten* KO), and Cre:ER^(T)+/-^Pten^F/F^miR-322^∼^503^-/-^ (dKO) mice.

To investigate the transcriptional impact of miR-322^∼^503 deletion, we first performed a linear multidimensional scaling analysis. This analysis revealed substantial variations in overall gene expression profiles across the different genotypes, while replicates within each genotype exhibited highly consistent patterns overall **(Figure 3A)**. Subsequently, we conducted a differential gene expression analysis (DEG, log_2_ FC > abs(1.5) plus adj. FDR p< 0.05) to compare gene expression profiles across the following groups: miR KO vs. Wt, *Pten* KO vs. Wt, dKO vs. Wt, and *Pten* KO vs. dKO **(Figure 3B-C**, **Supplementary Table S2)**. As anticipated, the comparison between *Pten* KO and Wt samples exhibited the highest number of genes with significantly altered expression (n=1,425), reflecting the profound transcriptional changes associated with PTEN loss. However, a notable number of DEGs were also identified in comparisons involving miR-322^∼^503-deficient samples, including miR KO vs. Wt (n=289) and dKO vs. *Pten* KO (n=210), underscoring the profound regulatory impact and gene expression dynamics influenced by miR-322^∼^503 deletion across experimental conditions. Then, to identify differentially regulated biological processes or molecular functions (i.e., pathway-level insights) underlying the reduced cell proliferation observed in organoids lacking miR-322^∼^503, we conducted an exploratory analysis by GSEA ^45^ using predefined gene set annotations. Given that the decrease in proliferation was evident in both Wt and *Pten* KO organoids deficient in miR-322^∼^503, we focused our analysis on comparing Wt and miR KO samples. This strategy was chosen to avoid confounding effects arising from *Pten* ablation, which could interfere with the specific impact of miR-322^∼^503 loss on gene expression. Our GSEA primarily retrieved gene signatures associated with proliferation and survival, including pathways regulating growth factor responses, insulin receptor signaling, and PI3K/AKT signaling **(Figure 3D)**, suggesting that the absence of miR-322^∼^503 disrupts the normal proliferation rate of endometrial epithelial cells by modulating these signaling cascades. These pathways are known to play critical roles in cellular growth and survival and are commonly dysregulated in endometrial epithelial cells. Among the identified pathways, two central signature hubs stood out due to their high statistical significance and relevance in explaining the effects of miR-322^∼^503 loss on endometrial cell proliferation. The first was the insulin-like growth factor 1 receptor (IGF1R)-coupled signaling pathway, a key regulator of cellular growth and metabolism. The second was the regulation of PI3K signaling, a central node in the control of cell proliferation and survival **(Figure 3E)**. Both pathways were significantly modulated in miR KO samples, highlighting their potential role as mechanisms by which miR-322^∼^503 deficiency suppresses the proliferation of endometrial glands. Interestingly, in light of the negative regulation of cell proliferation in miR-322^∼^503 knockout organoids, we further explored transcriptomic signatures related to the suppression of cellular growth. As expected, we observed a positive association between multiple signatures related to the negative regulation of cell growth and the absence of miR-322^∼^503 **(Figure 3F)**. These findings suggest that the loss of miR-322^∼^503 triggers a transcriptional program that actively suppresses cell proliferation, likely through its impact on growth factor signaling and downstream regulatory pathways.

**Figure 3.**
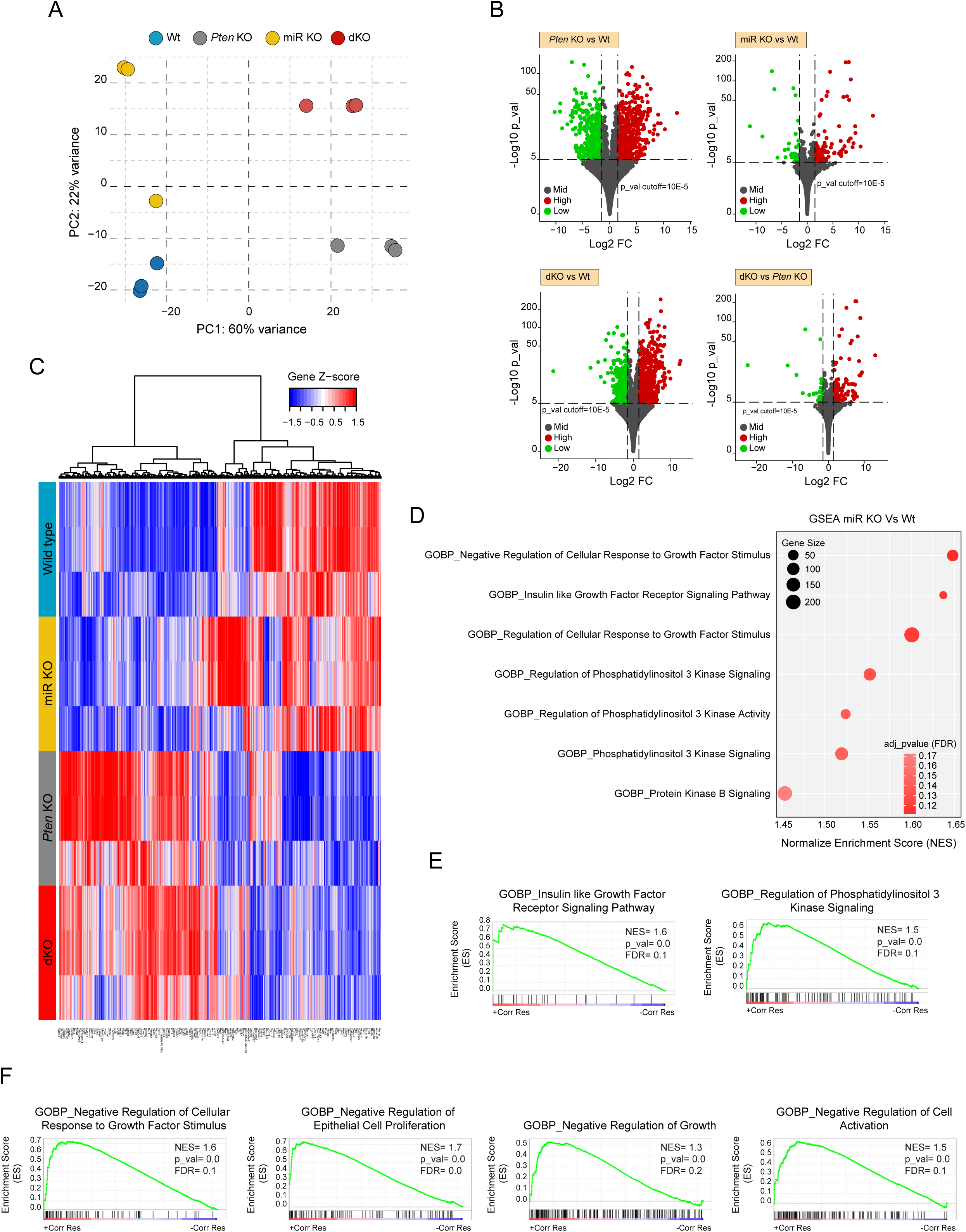
Loss of miR-322^∼^503 negatively regulates the transcriptome associated with growth factors and PI3K/AKT signaling. **(A)** Principal component analysis (PCA) of replicates from Cre:ER^(T)-/-^Pten^F/F^miR-322^∼^503^+/+^ (Wt), Cre:ER^(T)-/-^Pten^F/F^miR-322^∼^503^-/-^ (miR KO), Cre:ER^(T)+/-^ Pten^F/F^miR-322^∼^503^+/+^ (*Pten* KO), and Cre:ER^(T)+/-^Pten^F/F^miR-322^∼^503^-/-^ (dKO) samples. **(B)** Volcano plots of differentially expressed genes (DEGs) comparing the indicated genotypes. Green dots represent significantly upregulated genes, while red dots indicate significantly downregulated genes. **(C)** Heatmap of hierarchical clustering analysis showing DEGs across genotypes. **(D)** Dot plot illustrating gene set enrichment analysis (GSEA) of enriched gene signatures from the molecular signatures database (MSigDB) gene ontology (GO) biological process (BP). The plot highlights transcriptomic signatures related to cell growth and PI3K/AKT signaling in organoid cultures lacking miR-322^∼^503. Each node represents a distinct annotation with positive enrichment; node size corresponds to the number of genes included within the annotation. **(E)** Enriched gene annotations from GSEA (GO_BP) for "IGF1R receptor signaling pathway" and "PI3K signaling regulation." **(F)** GSEA plots of annotations associated with the negative regulation of cell proliferation and growth. RNA sequencing was performed on experimental samples (n=3), with each replicate derived from at least two animals per condition.

### miR-322^∼^503 knockout endometrial organoids display decreased Insulin/IGF-1 and PI3K/AKT signaling

To elucidate the role of miR-322^∼^503 in *Pten*-deficient endometrial organoids, we conducted a detailed analysis of the PI3K/AKT pathway, which is a critical regulator of endometrial epithelial cell proliferation and apoptosis in the context of *Pten* deficiency ^38^ **(Figure 4A)**. This pathway is known to be hyperactivated in Pten-deficient cells, contributing to their proliferative and survival advantages.

**Figure 4.**
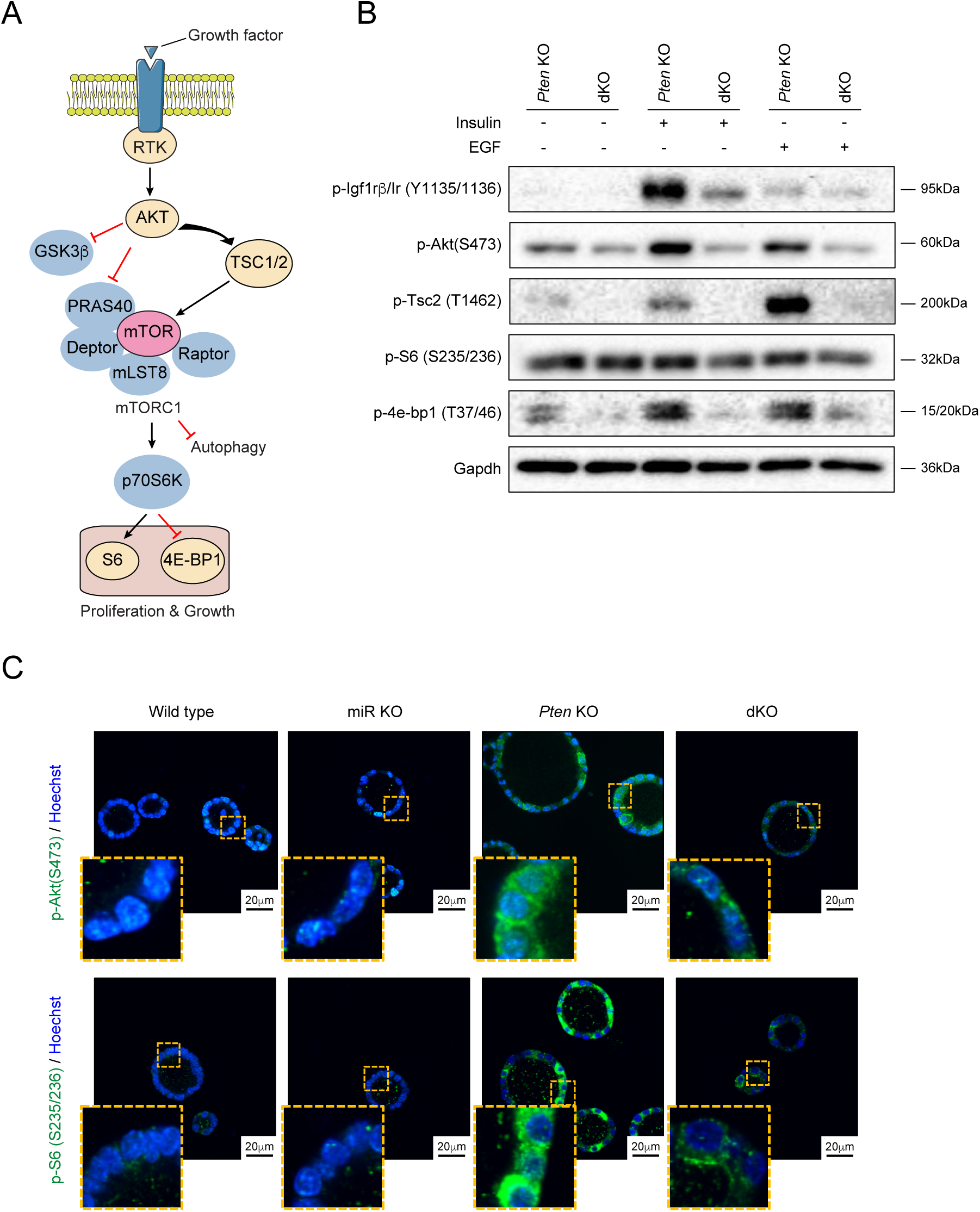
Loss of miR-322^∼^503 impairs insulin and EGF-induced activation of PI3K/AKT signaling. **(A)** Schematic diagram of receptor tyrosine kinase (RTK)-induced PI3K/AKT signaling. Nodes in yellow represent proteins analyzed in (B). **(B)** Representative western blot images of phosphorylated signaling proteins: p-Igf1rβ/Ir^Tyr1135/1136^, p-Akt^Ser473^, p-Tsc2^Thr1462^, p-S6^Ser235/236^ and p-4e-bp1^Thr37/46^. The analysis was performed on Cre:ER^(T)+/-^Pten^F/F^miR-322^∼^503^+/+^ (*Pten* KO), and Cre:ER^(T)+/-^Pten^F/F^miR-322^∼^503^-/-^ (dKO) organoid cultures stimulated with 1:100 dilution of Insulin-Transferrin-sodium selenite Supplement (ITS) or 5ηg/µL EGF for 10 minutes. Membranes were re-probed with Gapdh antibody as a protein loading control. **(C)** Representative confocal images showing immunofluorescence staining for p-AKT^Ser473^ and pS6^Ser235/236^ in organoids from Cre:ER^(T)-/-^Pten^F/F^miR-322^∼^503^+/+^ (Wt), Cre:ER^(T)-/-^Pten^F/F^miR-322^∼^503^-/-^ (miR KO), Cre:ER^(T)+/-^ Pten^F/F^miR-322^∼^503^+/+^ (*Pten* KO), and Cre:ER^(T)+/-^Pten^F/F^miR-322^∼^503^-/-^ (dKO) transgenic mice. Nuclei were stained with Hoechst for visualization. Scale bar: 20μm.

Of note, our organoid cultures grow in a defined medium that contains only EGF and insulin as growth factors, ensuring a controlled environment to investigate pathway activation ^37^. Consequently, PI3K/AKT signaling in these organoids is expected to depend solely on the presence of insulin and/or EGF. To assess the impact of miR-322^∼^503 loss on this pathway, we utilized mouse endometrial organoids with the following genotypes: Cre:ER^(T)+/-^Pten^F/F^miR-322^∼^503^+/+^ (*Pten* KO), and Cre:ER^(T)+/-^Pten^F/F^miR-322^∼^503^-/-^ (dKO). These organoids were stimulated with insulin and EGF, the only growth factors present in the culture medium, and pathway activation was analyzed via western blot. Western blot analysis revealed a pronounced reduction in the phosphorylation of key PI3K/AKT signaling components, including the IGF1/insulin receptor, AKT, TSC2, and 4E-BP1, in dKO organoids compared to *Pten* KO organoids. This reduction was observed under both basal (non-stimulated) and stimulated conditions **(Figure 4B)**. These findings indicate that miR-322^∼^503 deficiency significantly impairs the activation of the PI3K/AKT pathway in *Pten*-deficient cells. However, basal phosphorylation levels of PI3K/AKT in wild type (Wt) organoids were relatively low, and the limited protein yield from small organoids, particularly those with the miR-322^∼^503-deficient genotype, posed challenges for direct comparisons between Wt and miRNA KO organoids. To overcome these limitations and gain cellular resolution, we performed immunofluorescence analysis on organoids from Cre:ER^(T)-/-^Pten^F/F^miR-322^∼^503^+/+^ (Wt), Cre:ER^(T)-^

^/-^Pten^F/F^miR-322^∼^503^-/-^ (miR KO), Cre:ER^(T)+/-^Pten^F/F^miR-322^∼^503^+/+^ (*Pten* KO), and Cre:ER^(T)+/-^ Pten^F/F^miR-322^∼^503^-/-^ (dKO) mice. Immunofluorescence staining for phosphorylated AKT (p-AKT) and phosphorylated S6 (p-S6) corroborated the western blot findings. Specifically, dKO organoids exhibited markedly reduced phosphorylation levels of both AKT and S6 compared to *Pten* KO organoids. Similarly, miR KO organoids displayed decreased phosphorylation of these proteins relative to their Wt counterparts **(Figure 4C)**. These results collectively demonstrate that the absence of miR-322^∼^503 disrupts PI3K/AKT pathway activation in both *Pten*-proficient and *Pten*-deficient endometrial cells. This impairment underscores the critical role of miR-322^∼^503 in modulating signaling pathways essential for cell proliferation and survival in the context of *Pten* deficiency.

### Lack of miR-322^∼^503 restores TGF**b**-induced apoptosis in *Pten*-deficient endometrial organoids

We have previously demonstrated that *Pten* deficiency renders endometrial epithelial cells resistant to TGFβ-induced apoptosis through a PI3K/AKT-dependent mechanism ^38^. Building on this finding, we sought to investigate the potential involvement of miR-322^∼^503 in modulating apoptosis resistance to TGFβ in *Pten*-deficient organoids. Notably, miR-322^∼^503 has been shown to be activated by both TGFβ stimulation and PI3K/AKT signaling upregulation. Furthermore, our data indicate that the loss of miR-322^∼^503 diminishes PI3K/AKT pathway activation. To explore the role of miR-322^∼^503 in this context, we leveraged our transcriptomic analyses performed by GSEA to identify processes related to TGFβ signaling and apoptosis. This analysis revealed significant enrichment of several apoptosis and TGFβ-related processes that were impacted by miR-322^∼^503 deficiency **(Figure 5A-B)**. These findings suggest that miR-322^∼^503 plays a critical role in regulating the interplay between TGFβ signaling and apoptosis in *Pten*-deficient endometrial organoids.

**Figure 5.**
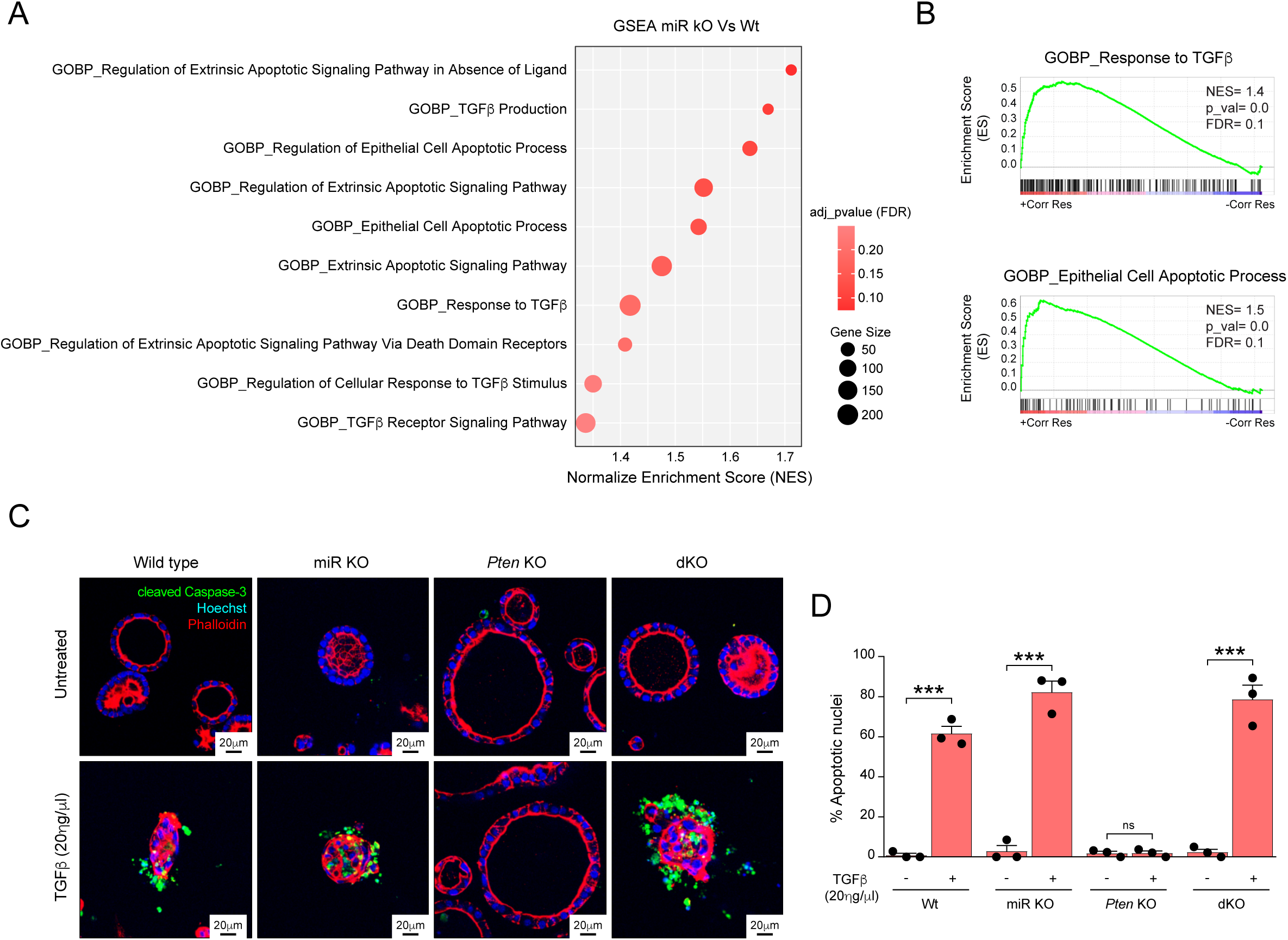
miR-322^∼^503 deficiency restores TGFβ-induced apoptosis in *Pten*-deficient organoids. **(A)** Dot plot illustrating gene set enrichment analysis (GSEA) of enriched gene signatures associated with apoptosis and TGFβ signaling in organoid cultures deficient in miR-322^∼^503. **(B)** Specific enriched gene annotations from GSEA (GO_BP) for "Response to TGFβ" and "Epithelial apoptotic cell process". **(C)** Representative confocal images of double immunofluorescence staining for cleaved caspase-3 and phalloidin in organoids from Cre:ER^(T)-/-^ Pten^F/F^miR-322^∼^503^+/+^ (Wt), Cre:ER^(T)-/-^Pten^F/F^miR-322^∼^503^-/-^ (miR KO), Cre:ER^(T)+/-^Pten^F/F^miR-322^∼^503^+/+^ (*Pten* KO), and Cre:ER^(T)+/-^Pten^F/F^miR-322^∼^503^-/-^ (dKO) mice. Organoids were stimulated with 20ηg/μL TGFβ for 48 hours. Phalloidin is visualized in red, cleaved caspase-3 in green, and nuclei are stained with Hoechst (blue). Scale bar: 20μm (60X magnification). **(D)** Quantification of apoptotic nuclei in organoids. Data are presented as mean ±SD, by t-test analysis, ***p<0.001; ns, not significant.

On the basis of these lines of evidence, we resolved to investigate whether the absence of miR-322^∼^503 influences the response of endometrial organoids to TGF-β treatment. To address this, Cre:ER^(T)-/-^Pten^F/F^miR-322^∼^503^+/+^ (Wt), Cre:ER^(T)-/-^Pten^F/F^miR-322^∼^503^-/-^ (miR KO), Cre:ER^(T)+/-^ Pten^F/F^miR-322^∼^503^+/+^ (*Pten* KO), and Cre:ER^(T)+/-^Pten^F/F^miR-322^∼^503^-/-^ (dKO) mouse-derived organoids were treated with 20ηg/µl TGFβ for 48 hours, followed by double immunofluorescence staining for cleaved caspase-3 to identify apoptotic nuclei and phalloidin to visualize actin filaments for enhanced cell visualization. As anticipated, Wt organoids exhibited robust activation of caspase-3, while *Pten* deficiency conferred complete resistance to TGFβ-induced apoptosis **(Figure 5C-D)**. Remarkably, TGFβ treatment led to a significant increase in apoptotic nuclei in dKO organoids compared to their *Pten* KO counterparts **(Figure 5C-D)**. In this line, the number of apoptotic nuclei in dKO organoids was comparable to that observed in Wt organoids, strongly supporting the critical role of miR-322^∼^503 in mediating apoptosis resistance associated with *Pten* deficiency.

### miR-322^∼^503 deficiency reduces tumor progression of endometrial cancer initiated by *Pten* loss *in vivo*

To investigate the role of miR-322^∼^503 deficiency in endometrial neoplasia, we employed our inducible *Pten* knockout mouse model. Importantly, the use of our Pten^F/F^ mice expressing a tamoxifen-inducible Cre-ER^(T)^ under the control of the chicken *Actb* promoter (CAGG) will circumvent the spontaneous development of aggressive neoplasms in non-target tissues, such as lymphoid tissues, which are commonly observed in other transgenic models like Pten^+/-^ heterozygous mice ^4 5^. Such off-target tumor development could interfere with the proper assessment of endometrial-specific pathology. By contrast, crossing Pten^F/F^ mice with CAGG-Cre:ER^(T)M^ will result in a model (Cre:ER^(T)+/-^Pten^F/F^) in which *Pten* recombination and deletion will occur specifically in epithelial tissues, including the endometrium, without affecting hematopoietic tissues ^36^. More specifically, after injection with a single dose of 4-tetrahydrotamoxifen (4-OHT), Cre:ER^(T)+/-^Pten^F/F^ will develop endometrial tumors within approximately 6 to 12 weeks. Hence, to further explore the interaction between PTEN loss and miR-322^∼^503 deficiency, we used four genotypes of 5- to 8-week-old female mice: Cre:ER^(T)-/-^ Pten^F/F^miR-322^∼^503^+/+^ (Wt), Cre:ER^(T)-/-^Pten^F/F^miR-322^∼^503^-/-^ (miR KO), Cre:ER^(T)+/-^Pten^F/F^miR-322^∼^503^+/+^ (*Pten* KO), and Cre:ER^(T)+/-^Pten^F/F^miR-322^∼^503^-/-^ (dKO). All animals received a single intraperitoneal injection of tamoxifen (0.5 mg/kg) after which they were monitored for up to 12 weeks post-injection. Euthanasia was performed as required and uteri were collected for histopathological analysis to assess tissue complexity and tumor progression **(Figure 6A)**. No significant correlation was observed between histopathological outcomes and euthanasia time points (*data not shown*); thus, all animals were analyzed as a single cohort. Next, anatomopathological evaluation of resected tissues revealed that miR-322^∼^503 deficiency significantly reduced tumor aggressiveness and, in some cases, prevented tumor formation in *Pten*-deficient endometrial tissue **(Figure 6B)**. Notably, 100% of Wt and miRNA KO mice exhibited no endometrial lesions. In contrast, all *Pten* KO mice developed endometrial neoplasia, with 10% presenting endometrial intraepithelial neoplasia (EIN) of complexity 1, 70% with EIN of complexity 2, and 20% with EIN of complexity 3. Among dKO mice, 31.25% showed no endometrial lesions, 21.87% exhibited complexity 1 EIN, 37.5% displayed complexity 2 EIN, and only 9.37% developed highly complex lesions **(Figure 6C)**. These findings underscore the critical role of miR-322^∼^503 in modulating tumor aggressiveness in *Pten*-deficient endometrial cancer.

**Figure 6.**
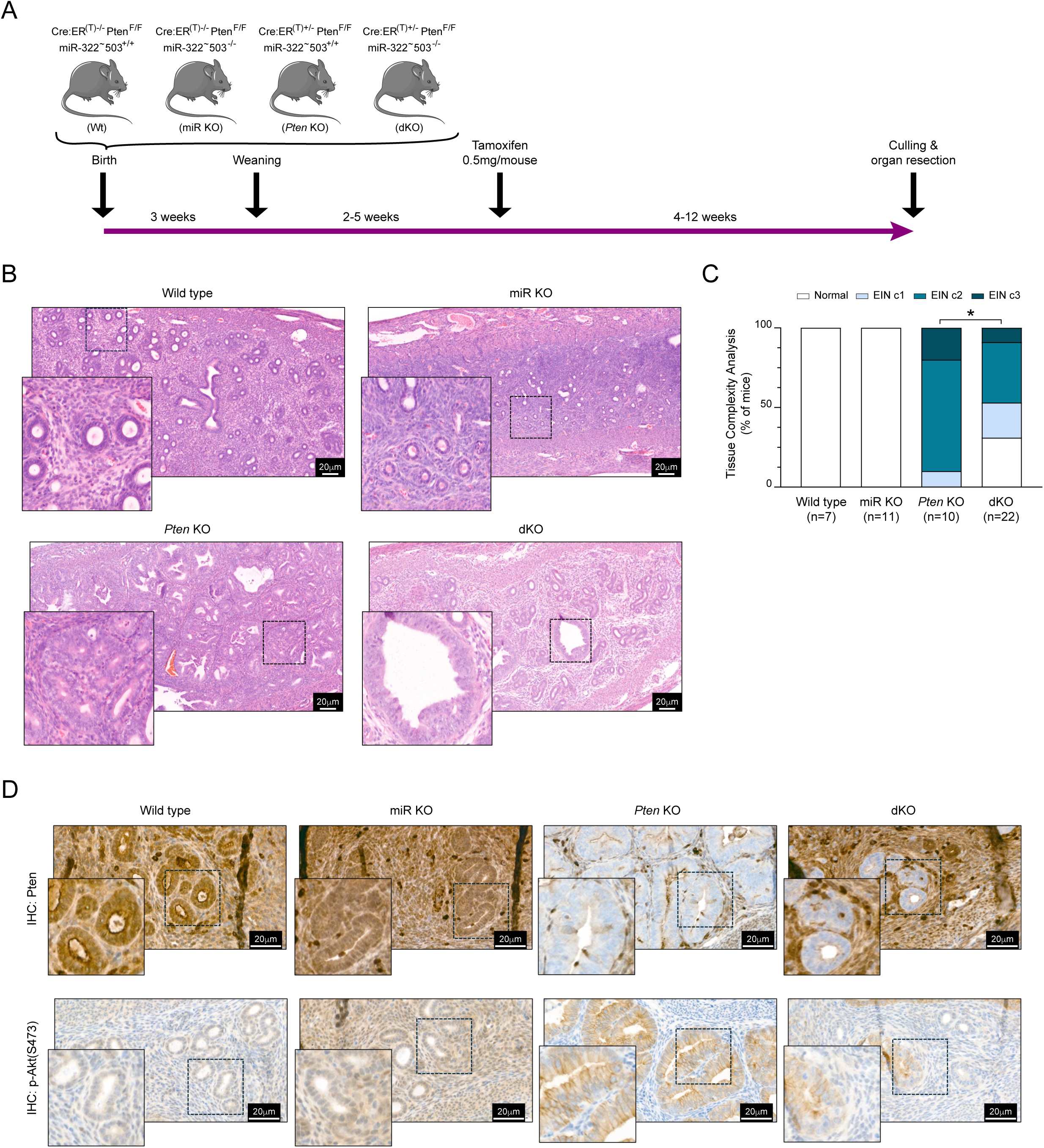
miR-322^∼^503 deficiency reduces tumor complexity in endometrial cancer initiated by *Pten* loss *in vivo*. **(A)** Timeline for the *in vivo* analysis of endometrial lesions in Cre:ER^(T)-/-^ Pten^F/F^miR-322^∼^503^+/+^ (Wt), Cre:ER^(T)-/-^Pten^F/F^miR-322^∼^503^-/-^ (miR KO), Cre:ER^(T)+/-^Pten^F/F^miR-322^∼^503^+/+^ (*Pten* KO), and Cre:ER^(T)+/-^Pten^F/F^miR-322^∼^503^-/-^ (dKO) mice **(B)** Representative hematoxylin and eosin (H&E) staining of endometrial sections from Wt, miR KO, *Pten* KO, and dKO mice. Scale bar: 20μm (40X magnification). **(C)** Histopathological analysis of endometrial sections shown in (B). Statistical significance was determined using Chi-squared analysis *p<0.05. **(D)** Representative immunohistochemistry images of p-Akt^Ser473^ and Pten in endometrial sections from WT, miR KO, *Pten* KO, and dKO mice. Scale bar: 20μm (40X magnification).

To ensure that the observed tumor shrinkage was not attributable to inefficient *Pten* ablation by tamoxifen, we performed immunohistochemical analysis of Pten protein expression in endometrial tissues across all four genotypes. Interestingly, while fewer glands exhibited negative staining for Pten in the dKO mice, those glands that had lost Pten in the dKO condition displayed a normal or less aggressive architecture compared to the glands observed in the *Pten* KO control group **(Figure 6D)**. To further investigate the downstream effects of *Pten* loss, we analyzed the activation of the PI3K/AKT signaling pathway by assessing phosphorylated Akt (p-Akt^Ser473^) levels via immunohistochemistry. In the dKO mice, *Pten*-deficient glands demonstrated a markedly lower intensity of Akt phosphorylation compared to glands from *Pten* KO mice **(Figure 6D)**. This suggests that miR-322^∼^503 deficiency mitigates the hyperactivation of the PI3K/AKT pathway typically associated with *Pten* loss. In the Wt and miR KO endometrial tissues, where basal PI3K/AKT activity is inherently low, no discernible differences in Akt phosphorylation were observed between the genotypes. These findings indicate that miR-322^∼^503 plays a significant role in modulating PI3K/AKT pathway activation in the context of *Pten* deficiency, fully contributing to the observed reduction in tumor aggressiveness.

### Double deficient *Pten* and miR-322^∼^503 uterine xenotransplants do not develop endometrial carcinomas

The tumor microenvironment (TME), composed of vascular cells, immune cells, and stromal cells, plays a critical role in cancer progression by influencing tumor growth, invasion, and metastasis. While our previous studies have demonstrated cell-intrinsic functions of miR-322^∼^503 in epithelial endometrial cells using organoid models, we sought to investigate whether the *in vivo* effects of miR-322^∼^503 deficiency might be mediated through stromal or immune cells and their interactions with epithelial cells. To address this question, we employed a dual stroma-epithelium xenograft model in severe combined immunodeficient (SCID) mice **(Figure 7A)**. Epithelial endometrial cells were isolated and cultured from Cre:ER^(T)-/-^Pten^F/F^miR-322^∼^503^+/+^ (Wt), Cre:ER^(T)-/-^Pten^F/F^miR-322^∼^503^-/-^ (miR KO), Cre:ER^(T)+/-^Pten^F/F^miR-322^∼^503^+/+^ (*Pten* KO), and Cre:ER^(T)+/-^Pten^F/F^miR-322^∼^503^-/-^ (dKO) trasgenic mice. In parallel, stromal and myometrial uterine cells were cultured from Wt mice. A 1:1 mixture of epithelial and stromal/myometrial cells was then subcutaneously inoculated into SCID mice. Ten weeks post-transplantation, animals were sacrificed, and subcutaneous tumors were harvested for macroscopic and histopathological analyses. Tumor volumes were calculated for xenografts derived from Wt stromal cells combined with epithelial cells of each genotype (Wt, miR KO, *Pten* KO, and dKO). As anticipated, tumor growth was observed only in xenografts containing *Pten*-deficient epithelial cells. Notably, epithelial cells lacking *Pten*, when combined with Wt stromal cells, produced significantly larger xenografts compared to all other conditions **(Figure 7B-C)**. Intriguingly, xenografts derived from dKO epithelial cells combined with Wt stromal cells were comparable in size to those generated with Wt epithelial cells, suggesting that miR-322^∼^503 deficiency impairs tumor progression of endometrial epithelial cells in a cell-autonomous manner. Histological analyses further supported these findings. In xenografts containing Wt or miR KO epithelial cells, tumors exhibited normal glandular structures surrounded by stromal cells, with epithelial cells displaying positive Pten staining **(Figure 7D)**. In contrast, xenografts with *Pten*-deficient epithelial cells (*Pten* KO) showed carcinomatous structures composed of *Pten*-deficient epithelial cells and Pten positive stromal cells **(Figure 7D**. In the dKO condition, xenografts displayed disorganized structures lacking *Pten* but showed no evidence of malignancy. These observations further underscore the critical role of miR-322^∼^503 in modulating tumor progression through cell-intrinsic mechanisms within endometrial epithelial cells, while its deficiency in the TME appears to have minimal impact on tumorigenesis.

**Figure 7.**
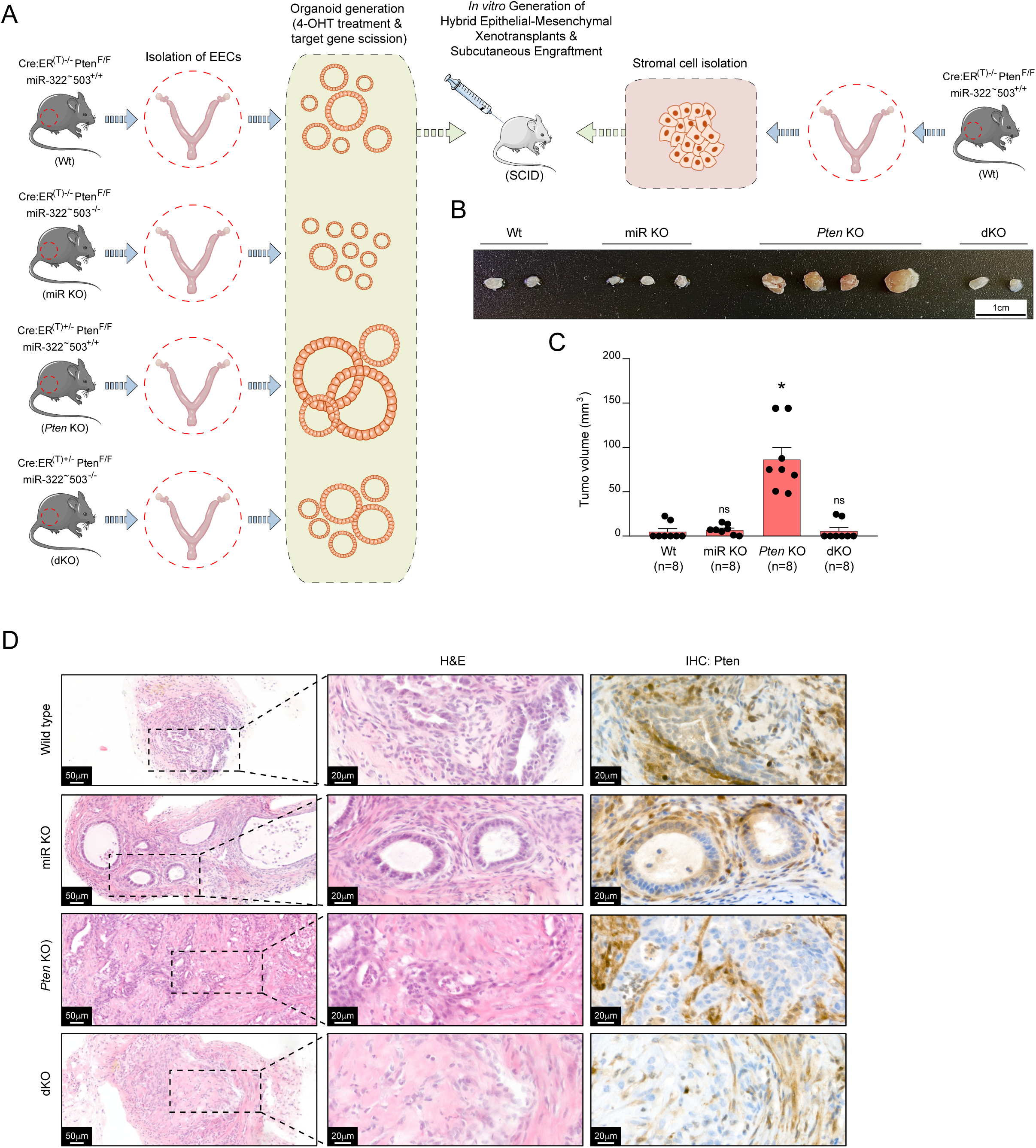
Endometrial xenotransplants containing Wt stromal cells and *Pten* and miR-322^∼^503 double-deficient endometrial epithelial cells do not exhibit endometrial intraepithelial neoplasia. **(A)** Schematic illustration of the experimental design. **(B)** Representative macroscopic images of subcutaneous xenografts derived from four combinations of a 1:1 mix of wild type stromal cells with epithelial endometrial cells from Cre:ER^(T)-/-^Pten^F/F^miR-322^∼^503^+/+^ (Wt), Cre:ER^(T)-/-^Pten^F/F^miR-322^∼^503^-/-^ (miR KO), Cre:ER^(T)+/-^Pten^F/F^miR-322^∼^503^+/+^ (*Pten* KO), and Cre:ER^(T)+/-^Pten^F/F^miR-322^∼^503^-/-^ (dKO) mice. Scale bar 1cm. **(C)** Volume quantification (mm^3^) of hybrid tumors formed by the combinations of Wt stromal cells with Wt, miR KO, *Pten* KO, and dKO epithelial cells. **(D)** Representative hematoxylin and eosin (H&E) staining and immunohistochemistry for Pten and p-Akt^Ser473^ in xenografts containing Wt stromal cells and Wt, miR KO, *Pten* KO, or dKO epithelial cells. Scale bar: 20μm (40X magnification). Data are presented as mean ±SD, by t-test analysis, *p<0.05; ns, not significant.

## DISCUSSION

MicroRNAs (miRNAs) are critical regulators of gene expression and have been implicated in various aspects of cancer biology, including tumor initiation, progression, and metastasis ^46^. In endometrial cancer (EC), while several studies have identified and validated the role of specific miRNAs, the functional characterization of many miRNAs remains incomplete. Notably, there is a lack of *in vivo* studies investigating the role of the miR-424(322)^∼^503 cluster in the endometrium, particularly using genetically engineered mouse models. In this study, we provide compelling evidence that the miR-424(322)^∼^503 cluster plays a pivotal role in the progression of EC driven by PTEN deficiency. Utilizing miR-424(322)^∼^503 knockout mice, we demonstrate that the absence of this miRNA cluster significantly impairs the development of EC initiated by conditional *Pten* loss. These findings suggest that the miR-424(322)^∼^503 cluster exhibits oncogenic properties in the context of the endometrium, promoting tumorigenesis in the setting of PTEN deficiency. Mechanistically, we reveal that the loss of miR-424(322)^∼^503 disrupts two key signaling pathways essential for endometrial tissue homeostasis: the PI3K/AKT pathway, which is frequently dysregulated in EC, and the TGFβ/Smad pathway, a critical mediator of apoptosis and cellular differentiation. These results underscore the multifaceted role of the miR-424(322)^∼^503 cluster in modulating signaling networks that are vital for maintaining endometrial integrity and highlight its potential as a therapeutic target in PTEN-deficient EC.

The regulation of miR-424(322)^∼^503 expression remains a complex and poorly understood process, involving multiple signaling pathways and transcriptional regulators. Emerging evidence highlights the intricate interplay between these factors in modulating miRNA expression in various physiological and pathological contexts. In our study, we identified a link between PTEN deficiency and the upregulation of miR-322 and miR-503 expression, a phenomenon that mirrors the effect of TGFβ treatment. TGFβ is known to induce miR-424(322)^∼^503 expression through the activation of Smad transcription factors, as we have previously reported ^23^ ^24^. Consistent with this mechanism, we found that the upregulation of miR-322 and miR-503 in *Pten*-deficient cells was partially attenuated by the addition of a PI3K inhibitor and completely abolished in the absence of Smad2/3. These findings align with prior work from our laborator, which demonstrated that *Pten* loss leads to constitutive nuclear translocation of Smad2/3 in the mouse endometrium ^42^. This nuclear localization likely facilitates Smad2/3 binding to the miR-424(322)/503 promoter, thereby driving its transcriptional activation. Collectively, these results suggest a model in which PTEN deficiency promotes miR-424(322)^∼^503 upregulation through a dual mechanism involving PI3K/AKT signaling and Smad2/3-dependent transcriptional regulation. This highlights the critical role of PTEN in maintaining miRNA homeostasis and underscores the importance of SMAD2/3 as a key mediator of miR-424(322)^∼^503 expression in the endometrium.

The observed upregulation of miR-424(322)^∼^503 expression prompted us to investigate its role in PTEN-loss-driven endometrial carcinogenesis, specifically whether it functions as a tumor promoter or suppressor. The role of the miR-424(322)^∼^503 cluster in cancer is highly context-dependent, with evidence supporting its dual functions as both an oncogene and a tumor suppressor across different cancer types. In various malignancies, including breast cancer, this cluster has been shown to exert opposing effects depending on the cellular and molecular context ^20-22^.

In endometrial cancer (EC), the role of miR-424(322)^∼^503 remains controversial. Some studies suggest that miR-424 acts as a tumor suppressor by targeting multiple oncogenic pathways. For instance, downregulation of miR-424 in human EC cell lines has been reported to inhibit metastasis by modulating the PTEN/PI3K/AKT signaling pathway ^35^. Additional studies have identified a tumor-suppressive role for miR-424 in EC cells, with targets including MMSET ^28^, E2F7 ^29^, E2F6 ^30^, PDIA6 ^31^, IGF-1R ^32^ and SPTBN2 ^33^. However, these investigations were primarily conducted *in vitro* using EC cell lines harboring diverse molecular alterations, which could influence the functional role of miR-424. Despite these insights, there are no *in vivo* studies addressing the role of miR-424(322)^∼^503 in EC using genetically engineered mouse models.

Our findings provide novel evidence that genetic ablation of miR-424(322)^∼^503 reduces EC development *in vivo* by attenuating cell proliferation and restoring TGFβ-induced apoptosis in *Pten*-deficient cells. Interestingly, the role of miR-424(322)^∼^503 in the endometrium may extend beyond tumorigenesis. Our results also reveal that knockout of miR-424(322)^∼^503 decreases proliferation in wild type endometrial epithelial cells, suggesting that this miRNA cluster may regulate broader aspects of endometrial biology. These findings raise the possibility that miR-424(322)^∼^503 contributes to other endometrial pathologies, such as endometriosis, as suggested by prior studies ^47-49^. Collectively, this highlights the multifaceted role of miR-424(322)^∼^503 in both normal and pathological endometrial physiology, emphasizing the need for further investigation into its molecular functions and therapeutic potential.

Mechanistically, our findings demonstrate that miR-424(322)^∼^503 knockout attenuates PI3K/AKT signaling, resulting in reduced endometrial cell proliferation and restoration of TGFβ-induced apoptosis. This observation contrasts with studies in mammary epithelium, where miR-424(322)^∼^503 targeted deletion in mice has been shown to exert tumor-suppressive effects by targeting key regulators such as CDC25A, BCL-2, and IGF-1R ^23^ ^24^ ^50^ and to promote metaplastic differentiation and stem cell expansion by regulating LRP6 and β-catenin/WNT signaling ^51^. However, our transcriptomic analysis did not identify significant changes in the expression of these targets in miR-424(322)^∼^503-deficient endometrial cells. Instead, we observed transcriptional signatures indicative of reduced cell growth, proliferation, and responsiveness to growth factors, particularly those associated with Insulin/EGF signaling. These findings aligns with the culture conditions of our organoid models, which utilize a defined medium containing EGF and insulin as the sole growth factors. Both are essential for the development and maintenance of endometrial organoids ^37^. Consequently, the observed decrease in Insulin/EGF signaling, coupled with diminished PI3K/AKT activation, provides a plausible explanation for the reduced proliferation seen in miR-424(322)^∼^503-deficient cells. Moreover, previous work from our laboratory has established that PTEN loss alone is sufficient to drive excessive endometrial cell proliferation and to inhibit TGFβ-induced apoptosis via a PI3K/AKT-dependent mechanism ^38^. In this context, the attenuation of PI3K/AKT signaling caused by miR-424(322)^∼^503 knockout in *Pten*-deficient cells is particularly noteworthy, as it not only reduces proliferation but also restores the apoptotic response to TGFβ. These findings underscore the tissue-specific nature of miR-424(322)^∼^503 function, which may explain its contrasting roles in cancer in different cellular contexts. In the endometrium, miR-424(322)^∼^503 fine-tunes key signaling pathways essential for cellular proliferation and survival, emphasizing its importance in PTEN-loss-driven endometrial carcinogenesis.

One limitation of the mouse model utilized in this study is that it involves a complete knockout of the miR-424(322)^∼^503 cluster. Consequently, the observed *in vivo* effects may not be exclusively attributable to the loss of miR-424(322)^∼^503 in epithelial endometrial cells. Instead, the deletion of this miRNA cluster in non-epithelial cell populations, such as stromal, vascular, immune, or adipose cells, could also influence the development and progression of endometrial cancer (EC). This is particularly relevant given that miR-424(322)^∼^503 knockout has been implicated in other pathophysiological conditions, including obesity ^52^, a known risk factor that can impact cancer initiation and progression.

The potential contribution of non-epithelial cells to the observed phenotypes in *Pten*-deficient epithelial cells underscores the complexity of tumor-microenvironment interactions. For example, miR-424(322)^∼^503 deficiency in stromal or immune cells could alter cytokine secretion, extracellular matrix remodeling, or angiogenesis, indirectly affecting epithelial cell behavior. Despite these considerations, our *in vivo* xenograft experiments, in which *Pten*-deficient epithelial cells were combined with wild type stromal cells, strongly support a cell-autonomous role for miR-424(322)^∼^503 in endometrial epithelial cells. Furthermore, our *in vitro* organoid studies provide additional evidence that the ablation of miR-424(322)^∼^503 directly impairs epithelial cell proliferation and survival. These findings suggest that while the broader effects of miR-424(322)^∼^503 deficiency in non-epithelial cells cannot be entirely excluded, the primary driver of the observed phenotypes in our model is likely the intrinsic loss of miR-424(322)^∼^503 within the epithelial compartment. Future studies employing cell-type-specific knockouts of miR-424(322)^∼^503 will be essential to delineate its distinct roles in epithelial versus non-epithelial cells and to fully understand its contribution to the complex interplay between the tumor and its microenvironment.

## MATERIALS AND METHODS

### Experimental Mouse Models

Mice were housed in a barrier facility and pathogen-free procedures were used in all mouse rooms. Animals were under 12 hours of light/dark cycles at 22 °C, and they had *ad libitum* access to water and food. Procedures in this study were done following the guidelines of Ethical Committee of Universitat de Lleida and the National Institute of Health Guide for the Care and Use of Laboratory Animals. Conditional *Pten* knockout (C; 129S4-Ptentm1Hwu/J or *Pten*^F/F^) and Cre:ER^(T)^ mice were obtained from the Jackson Laboratory (Bar Harbor, ME, USA). MiR-322^∼^503^-/-^ (FVB/NJ) mice were a gift from Prof. Jose Silva. Cre:ER^(T)+/-^Pten^F/F^miR-322^∼^503^-/-^ mice were bred by crossing Cre:ER^T+/-^*Pten*^F/F^ and miR-322^∼^503^-/-^ mice. Three weeks after birth, animals were weaned and genotyped as previously described ^23^ ^36^ ^38^. Genotyping primers and PCR conditions are specified in Supplementary Materials (Table S3). Immunodefficient female SCID^hr/hr^ mice were bred at the Universitat de Lleida housing facility.

### Isolation of Epithelial Endometrial Cells and Organoid Culture

Isolation of mouse epithelial endometrial cells was done as previously described ^37^. Briefly, mice were euthanized by cervical dislocation and uteri were dissected. Uterine horns were cut into 3mm pieces and washed in Hanks Balanced Salt Solution (HBSS) (14175-046, Gibco). Uterine fragments were digested with 1% trypsin (15090-046, Gibco) in HBSS for 1 hour at 4°C and for 45 minutes at room temperature. DMEM (41965-039, Gibco) with 10 % of Fetal Bovine Serum (FBS) (A52567-01, Gibco) was added to stop trypsin reaction. Then, with the edge of a razor blade, epithelial sheets were squeezed out of the uterine fragments. Epithelial sheets were washed twice with Phosphate Buffered Saline (PBS) and resuspended in 1 ml of basal medium (DMEM F/12 (11039-021, Gibco) with 1mmol HEPES (H0887, Sigma-Aldrich), 1 % penicillin/streptomycin (15140-122, Gibco) and amphotericin B (15290-018, Gibco). Epithelial sheets were mechanically disrupted in clusters of cells by pipetting 50 times with a 1ml tip. Clusters were diluted in basal medium supplemented with 2% dextran-coated charcoal-stripped serum (DCC) (A33821-01, Gibco) and plated into culture dishes (BD Falcon). When indicated, cells were treated with 0.5mM (Z)-4-HydroxyTamoxifen (H7904 Sigma-Aldrich) to promote ablation of LoxP flanked genes. Cells were incubated for 24 hours in an incubator at 37°C and 5% CO_2_. Then, cells were washed with PBS and incubated with trypsin/EDTA solution (25200-056, Gibco) for 5 minutes at 37°C. DMEM 10% FBS was added to stop trypsin reaction and cells were washed with PBS and centrifuged at 1000 rpm for 3 minutes. Cell pellet was resuspended in basal medium containing 3% Matrigel™ (354234, Corning) to obtain 4 x 10^4^ cell clumps/ml and plated on top of a Matrigel layer for each experiment. For RNA or protein extraction, cells were seeded in 24-well plates in a volume of 300μl. For immunofluorescence, cells were plated in a 96-well plate (black with micro-clear bottom) (Greiner Bio-one) in a volume of 80μl. In both cases, after incubating cells at 37°C and 5% CO_2_ for 24 hours, medium was replaced by basal medium supplemented with 5ηg/ml EGF (E9644, Sigma-Aldrich) and 1:100 dilution of Insulin-Transferrin-sodium selenite Supplement (ITS) (41400-045, Gibco) and 3% of Matrigel. Every 2-3 days medium was replaced until organoids were completely formed. Organoid treatments were performed two days after medium replacement. TGF-β (#PHG9214, Gibco) treatments were performed at the indicated doses and times.

### Whole genome miRNA sequencing

TruSeq small RNA prep, 1x50bp single end reads, HiSeq 2000, was conducted at New York Genome Center. Adapter sequence were removed from FASTQ file with fastx_clipper and alignment of reads to NCBI genome build 37, as well as to miRBase hairpin sequences and other small RNA sequences (tRNA, snoRNA, rRNA, snRNA) were conducted using Bowtie ^53^. miRBase transcripts were quantified using in-house scripts implemented in Python and imported into R for analysis with the DESeq package. For differential miRNAseq expression analysis low-expression miRNAs, defined as those with fewer than 10 reads across all samples, were filtered out prior to analysis. Data normalization and differential expression analysis between *Pten* KO, and Wt were performed using the DESeq2 R package (v1.30.1) ^54^.

### Whole genome mRNA sequencing

2x150bp stranded mRNA sequencing was conducted at Centre Nacional d’Anàlisi Genòmica (CNAG) on a NovaSeq 6000 equipment at a depth coverage of 20M reads per sample. Quality control of all FASTQ files was conducted using the FASTQC tool (v0.11.9) and MultiQC tool ^55^ implemented in R. For mRNAseq quantification the mapping-based mode of Salmon (v1.5.2) ^56^ was used to perform transcript-level quantification with default parameters, using the GRCm38 reference genome. Transcript-level data were imported into R and summarized to gene-level counts using the tximport library (v1.28.0) ^57^. Low-expression genes, defined as those with fewer than 10 reads across all samples, were filtered out prior to analysis. Data normalization and differential expression analysis between miR KO, *Pten* KO, dKO, and WT conditions, as well as between dKO and *Pten* KO, were performed using the DESeq2 R package (v1.30.1) ^54^. To conduct Gene Set Enrichment Analysis (GSEA ^45^), genes from the miR KO vs WT comparison were ranked by fold change and analyzed using the Pre-ranked mode of the Broad Institute’s Gene Set Enrichment Analysis (GSEA v4.0.3) software. The analysis utilized GO BP mouse gene sets from the MsigDB database ^58^, accessed through the msigdbr R function. Selected pathways were visualized in a dot plot, displaying FDR values and gene set sizes, created using the ggplot2 package in R.

### Hybrid Epithelial-Mesenchymal Xenotransplants and Isolation of Mesenchymal and Epithelial Cells from Mouse Uterus

Hybrid epithelial-mesenchymal uterine xenotransplants were generated as previously described ^60^. Concisely, mice were euthanized by cervical dislocation and uteri were dissected. For mesenchymal cells, uterine horns were cut into 3mm pieces and washed in HBSS. Then, uterine fragments were digested with 1% trypsin in HBSS for 1 hour at 4°C and for 45 minutes at room temperature. DMEM with 10% of FBS was added to stop trypsin reaction. Then, with the edge of a razor blade, epithelial sheets were squeezed out of the uterine fragments and removed. Next, myometrial and stromal cells remaining in the uterine pieces were incubated in digestion medium (2mg/ml collagenase type I (LS0004196, Worthington) and 5% FBS in DMEM) during 3 hours at 37°C. Cells were passed through a 40μm strainer and washed twice with PBS. Then cells were centrifuged at 1000 rpm for 3 minutes, resuspended in stromal medium (DMEM/F12 supplemented with 1 mM HEPES, 1% penicillin/streptomycin, 0.1% amphotericin B and 2% DCC) and plated in culture dishes. Cells were incubated at 37°C and 5% CO_2_ for two days or until they were at high confluence. Epithelial endometrial cells for xenotransplants were isolated and cultured in a monolayer as explained before. Then, A 1:1 mixture of epithelial / mesenchymal cells was resuspended in PBS with stromal medium and 20% Matrigel™. Cells were subcutaneously injected in a volume of 100μl in each flank of SCID mice.

### Tamoxifen Administration

Tamoxifen (T5648, Sigma-Aldrich) was dissolved and administered as previously described ^36^. In brief, tamoxifen powder was dissolved in 100% ethanol at a 100mg/ml concentration. Tamoxifen was emulsified in corn oil (C8267, Sigma-Aldrich) at 10mg/ml by vortexing. Adult females of 5-week-old were given a single intraperitoneal injection of 0.5mg of tamoxifen emulsion (30 - 35μg/mg body weight).

### Glandular Perimeter Measurements

Images of mouse endometrial organoids were taken with a phase-contrast microscope (Eclipse Ts2R, Nikon). Spheroid perimeter analysis was performed in an image analyzer (Image J, version 1.46r; NIH, Bethesda, MD, USA) by generation of binary images of the glands, as previously described ^61^.

### miRNA Extraction and Real-Time qPCR

MicroRNA extraction from organoid cultures was performed with mirVana miRNA isolation kit (AM1561, Ambion) according to manufacturer’s protocol. MiRNA extracts were quantified with Nanodrop (ND-1000 UV/Vis Spectophotometer, Nanodrop Technologies) and stored frozen at -80°C. Retrotranscription was performed with 50ηg of RNA using High-Capacity cDNA Reverse Transcription (4368815, Applied Biosystems), following manufacturer’s instructions. Primers and PCR conditions for reverse transcription are detailed in Supplementary Materials (Table S4). Quantitative real-time PCR detection of miRNAs was performed with the CFX96 (Bio-Rad) using Power Up SYBR Green Master Mix (A25742, Applied Biosystems). The sequences of primers used are available in Supplementary Materials and were obtained from IDT. Relative expression was determined from cycle threshold (Ct) values, which were normalized to sno-202 as a housekeeping miRNA. Experiments were performed at least three times, and every treatment group was performed in triplicate.

### Total RNA Extraction, Real-Time qPCR

Total RNA from organoid cultures was extracted using SurePrep TrueTotal RNA Purification Kit (BP2800-50, Fisher BioReagents) according to manufacturer’s protocol. Quantitative real-time PCR was performed with 50ηg of total RNA using the one-step protocol qPCRBIO Probe 1-Step Go (PB25.44-01, PCR Biosystems). Primers used for gene expression analysis were commercially obtained from Applied Biosystems: *Ccnd1* (Mm00432359_m1) and *Gapdh* (Mm99999915_g1). Relative expression was determined by cycle threshold (Ct) values, which were normalized to Gapdh expression as an internal control. Experiments were performed at least three times.

### Western Blot

Western blot analysis was performed as previously described ^36^ with minor variations. To dissociate organoids from Matrigel, cell cultures were washed with PBS and incubated at 37 °C for 15 minutes with TrypLE Express (12604-013, Gibco) dissociation reagent. Cells were washed in PBS and centrifuged at 1000 rpm for 3 minutes. Cell pellet was lysed with lysis buffer (2% SDS, 125mM Tris-HCl pH 6.8). Relative protein concentrations were established with a colorimetric protein assay kit (5000112, BioRad). Equal amounts of protein were loaded to an acrylamide gel, transferred to PDVF membranes (IPVH00010, Millipore). Membranes were blocked for 1 h with TBS-T (20mM Tris-HCl pH 7.4, 150mM NaCl, 0.1% Tween-20) plus 5% of non-fat milk to avoid non-specific binding. Then, membranes were incubated overnight at 4°C with primary antibodies. Subsequently, membranes were incubated for 1 hour with secondary antibody at room temperature. Finally, signal was detected with Immobilon Forte Western HRP Substrate (WBLUF0100, Millipore). All primary and secondary antibodies used for Western blot and its concentrations are listed in Supplementary Materials (Table S5).

### Bromo-2’-deoxyuridine (BrdU) Labeling

Three-dimensional cultures were incubated with 3ng/ml of BrdU (B5002, Sigma-Aldrich) for 16 hours and were fixed with 4% paraformaldehyde (PFA) for 5 minutes. Denaturation of DNA was performed with 2M HCl for 30 minutes at 37°C. Then, HCl was neutralized with 0.1M of sodium tetraborate (pH 8.5) for 2 minutes and washed three times with PBS. Cells were blocked with BrdU blocking solution (5% horse serum, 5% FBS, 0.2% glycine and 0.1% Triton X-100) for 1 hour at room temperature. Then, cells were incubated overnight with a monoclonal antibody against BrdU and washed with PBS. Finally, cells were incubated overnight with anti-rat Alexa Fluor conjugated antibody and Hoechst as a nuclear marker. Endometrial glands were visualized under a confocal microscope (Olympus, Fluoview FV1000) and images were taken and analyzed with Olympus Fluoview software (FV19-AS4 version 4.2, Olympus). Positive BrdU cells were counted and divided by total cells (labelled with Hoechst) and results were expressed as BrdU-positive cells. Antibodies used for BrdU labelling are listed in Supplementary Materials (Table S4).

### Immunofluorescence

Immunofluorescence was done as previously described ^37^. In brief, endometrial organoids were fixed with 4% PFA in PBS for 5 minutes at room temperature, washed twice with PBS and permeabilized with 0.1% triton X-100 in PBS for 10 minutes and incubated in blocking solution (2% Horse Serum, 2% Bovine Serum Albumin, 0.2% Triton X-100 in PBS) for 1 hour at room temperature. Then, cells were incubated with the indicated dilution of primary antibodies overnight at 4°C. Cultures were washed twice with PBS and incubated in secondary Alexa Fluor conjugated antibodies and Hoechst, as a nuclear staining, overnight at 4°C. All antibodies and dilutions used for immunofluorescence are listed in Supplementary Materials (Table S6). Importantly, in double staining different isotypes of antibodies were used. Visualization and image capturing of immunofluorescence was performed with a confocal microscope (Olympus, Fluoview FV1000) and analyzed with Olympus Fluowiew software (FV19-AS4 version 4.2, Olympus).

### Tissue Processing and Immunohistochemistry Analysis on Paraffin Sections

For histological examination, animals were euthanized, and uteri were collected, formalin-fixed overnight at 4°C and paraffin-embedded. Paraffin sections of 3μm were dried for an hour at 80°C, dewaxed in xylene, gradually rehydrated in ethanol and washed in PBS. Antigen retrieval was performed in EnVision FLEX and high pH or low pH solution (K8004 or K8005, DAKO) for 20 minutes at 95°C, depending on each antibody. Then, endogenous peroxidase was blocked through 3% H_2_O_2_ incubation, and slides were washed three times with PBS. After that, primary antibodies were incubated for 30 minutes at room temperature, washed in PBS and incubated with Horseradish Peroxidase (HRP)-conjugated secondary antibodies. Finally, staining was visualized through reaction with EnVisioin Detection Kit (K4065, DAKO) using diaminobenzidine (DAB) substrate. Slides were counterstained with Harrys hematoxylin. Primary and secondary antibodies used for immunohistochemistry and their dilutions are listed in Supplementary Materials (Table S7).

### Statistical analysis

Statistical analysis was performed according to each experiment. Briefly, comparisons between two groups were analyzed with Student’s *t*-test and represented as mean ± standard deviation. Comparisons between two or more groups were done by two-way ANOVA. Finally, contingency tables were analyzed by χ^2^□test, followed by Fisher’s exact test. All experiments were at least performed three times, and all treatment groups were done in triplicates. GraphPad Prism (Version 8.0, GraphPad Software, Inc.) was used for statistical analysis.

## ACKNOWLEDGEMENTS

This study has been funded by the Institut de Recerca Biomédica de Lleida (PIRS2023, XD), Ministerio de Ciencia, Innovación y Universidades (PID2022-141220OB-I00, XD and PID2019-104734RB-I00, XD), and Instituto de Salud Carlos III (ISCIII) (PI21/00672, DL-N and PI24/01255, DL-N) (co-funded by the European Regional Development Fund. ERDF, a way to build Europe), and by the CIBERonc network (XM-G, CB16/1200231). We thank the Generalitat of Catalonia, Agency for Management of University and Research Grants (2021SGR01609 and 2021SGR01098). The authors also want to thank the CERCA programme / Generalitat de Catalunya for institutional support.

## AUTHOR CONTRIBUTIONS

MV-S, DL-N, and XD designed the research and developed the project concept. MV-S, DL-N, and XD prepared the manuscript. JE, ME, RR-B, JMS, XM-G, DL-N, and XD provided essential materials and resources necessary for conducting the research. MV-S, NB, RN, and XD performed the experiments and collected the data. MV-S, NB, RN, DL-N, and XD analyzed the data and conducted statistical analyses. All authors contributed to data interpretation and critically revised the manuscript.

## REFERENCES

1. Makker V, MacKay H, Ray-Coquard I, et al. Endometrial cancer. Nat Rev Dis Primers 2021;7(1):88.

2. Crosbie EJ, Kitson SJ, McAlpine JN, Mukhopadhyay A, Powell ME, Singh N. Endometrial cancer. Lancet 2022;399(10333):1412–28.

3. Di Cristofano A, Pesce B, Cordon-Cardo C, Pandolfi PP. Pten is essential for embryonic development and tumour suppression. Nat Genet 1998;19(4):348–55.

4. Podsypanina K, Ellenson LH, Nemes A, et al. Mutation of Pten/Mmac1 in mice causes neoplasia in multiple organ systems. Proc Natl Acad Sci U S A 1999;96(4):1563–8.

5. Suzuki A, de la Pompa JL, Stambolic V, et al. High cancer susceptibility and embryonic lethality associated with mutation of the PTEN tumor suppressor gene in mice. Curr Biol 1998;8(21):1169–78.

6. Navaridas R, Vidal-Sabanes M, Ruiz-Mitjana A, et al. Transient and DNA-free in vivo CRISPR/Cas9 genome editing for flexible modeling of endometrial carcinogenesis. Cancer Commun (Lond) 2023;43(5):620–24.

7. Maru Y, Hippo Y. Two-Way Development of the Genetic Model for Endometrial Tumorigenesis in Mice: Current and Future Perspectives. Front Genet 2021;12:798628.

8. Luo F, Huang Y, Li Y, et al. A narrative review of the relationship between TGF-beta signaling and gynecological malignant tumor. Ann Transl Med 2021;9(20):1601.

9. Zakrzewski PK. Canonical TGFbeta Signaling and Its Contribution to Endometrial Cancer Development and Progression-Underestimated Target of Anticancer Strategies. J Clin Med 2021;10(17).

10. Piestrzeniewicz-Ulanska D, Brys M, Semczuk A, Rechberger T, Jakowicki JA, Krajewska WM. TGF-beta signaling is disrupted in endometrioid-type endometrial carcinomas. Gynecol Oncol 2004;95(1):173–80.

11. Parekh TV, Gama P, Wen X, et al. Transforming growth factor beta signaling is disabled early in human endometrial carcinogenesis concomitant with loss of growth inhibition. Cancer Res 2002;62(10):2778–90.

12. Muinelo-Romay L, Colas E, Barbazan J, et al. High-risk endometrial carcinoma profiling identifies TGF-beta1 as a key factor in the initiation of tumor invasion. Mol Cancer Ther 2011;10(8):1357–66.

13. Mhawech-Fauceglia P, Kesterson J, Wang D, et al. Expression and clinical significance of the transforming growth factor-beta signalling pathway in endometrial cancer. Histopathology 2011;59(1):63–72.

14. Monsivais D, Peng J, Kang Y, Matzuk MM. Activin-like kinase 5 (ALK5) inactivation in the mouse uterus results in metastatic endometrial carcinoma. Proc Natl Acad Sci U S A 2019;116(9):3883–92.

15. Kriseman M, Monsivais D, Agno J, Masand RP, Creighton CJ, Matzuk MM. Uterine double-conditional inactivation of Smad2 and Smad3 in mice causes endometrial dysregulation, infertility, and uterine cancer. Proc Natl Acad Sci U S A 2019;116(9):3873–82.

16. Gao Y, Lin P, Lydon JP, Li Q. Conditional abrogation of transforming growth factor-beta receptor 1 in PTEN-inactivated endometrium promotes endometrial cancer progression in mice. J Pathol 2017;243(1):89–99.

17. Liu Q, Fu H, Sun F, et al. miR-16 family induces cell cycle arrest by regulating multiple cell cycle genes. Nucleic Acids Res 2008;36(16):5391–404.

18. Finnerty JR, Wang W-X, Hébert SS, Wilfred BR, Mao G, Nelson PT. The miR-15/107 Group of MicroRNA Genes: Evolutionary Biology, Cellular Functions, and Roles in Human Diseases. Journal of Molecular Biology 2010;402(3):491–509.

19. Wang F, Liang R, Tandon N, et al. H19X-encoded miR-424(322)/-503 cluster: emerging roles in cell differentiation, proliferation, plasticity and metabolism. Cellular and molecular life sciences : CMLS 2019;76(5):903–20.

20. Ghafouri-Fard S, Askari A, Hussen BM, Taheri M, Akbari Dilmaghani N. Role of miR-424 in the carcinogenesis. Clinical & translational oncology : official publication of the Federation of Spanish Oncology Societies and of the National Cancer Institute of Mexico 2024;26(1):16–38.

21. Li S, Wu Y, Zhang J, Sun H, Wang X. Role of miRNA-424 in Cancers. Onco Targets Ther 2020;13:9611–22.

22. Wu Y, Wang W, Yang AG, Zhang R. The microRNA-424/503 cluster: A master regulator of tumorigenesis and tumor progression with paradoxical roles in cancer. Cancer Lett 2020;494:58–72.

23. Llobet-Navas D, Rodriguez-Barrueco R, Castro V, et al. The miR-424(322)/503 cluster orchestrates remodeling of the epithelium in the involuting mammary gland. Genes Dev 2014;28(7):765–82.

24. Llobet-Navas D, Rodriguez-Barrueco R, de la Iglesia-Vicente J, et al. The microRNA 424/503 cluster reduces CDC25A expression during cell cycle arrest imposed by transforming growth factor beta in mammary epithelial cells. Mol Cell Biol 2014;34(23):4216–31.

25. Li Y, Li W, Ying Z, et al. Metastatic heterogeneity of breast cancer cells is associated with expression of a heterogeneous TGFbeta-activating miR424-503 gene cluster. Cancer Res 2014;74(21):6107–18.

26. Zhao Z, Fan X, Jiang L, et al. miR-503-3p promotes epithelial-mesenchymal transition in breast cancer by directly targeting SMAD2 and E-cadherin. J Genet Genomics 2017;44(2):75–84.

27. Wei S, Li Q, Li Z, Wang L, Zhang L, Xu Z. miR-424-5p promotes proliferation of gastric cancer by targeting Smad3 through TGF-beta signaling pathway. Oncotarget 2016;7(46):75185–96.

28. Dong P, Xiong Y, Yue J, Hanley SJB, Watari H. miR-34a, miR-424 and miR-513 inhibit MMSET expression to repress endometrial cancer cell invasion and sphere formation. Oncotarget 2018;9(33):23253-63.

29. Li Q, Qiu XM, Li QH, et al. MicroRNA-424 may function as a tumor suppressor in endometrial carcinoma cells by targeting E2F7. Oncol Rep 2015;33(5):2354–60.

30. Lu Z, Nian Z, Jingjing Z, Tao L, Quan L. MicroRNA-424/E2F6 feedback loop modulates cell invasion, migration and EMT in endometrial carcinoma. Oncotarget 2017;8(69):114281–91.

31. Wang P, Zhang T, Jiang N, et al. PDIA6, which is regulated by TRPM2-AS/miR-424-5p axis, promotes endometrial cancer progression via TGF-beta pathway. Cell Death Dis 2023;14(12):829.

32. Shu S, Liu X, Xu M, et al. MicroRNA-424 regulates epithelial-mesenchymal transition of endometrial carcinoma by directly targeting insulin-like growth factor 1 receptor. J Cell Biochem 2019;120(2):2171–79.

33. Wang P, Liu T, Zhao Z, Wang Z, Liu S, Yang X. SPTBN2 regulated by miR-424-5p promotes endometrial cancer progression via CLDN4/PI3K/AKT axis. Cell Death Discov 2021;7(1):382.

34. Devor EJ, Cha E, Warrier A, Miller MD, Gonzalez-Bosquet J, Leslie KK. The miR-503 cluster is coordinately under-expressed in endometrial endometrioid adenocarcinoma and targets many oncogenes, cell cycle genes, DNA repair genes and chemotherapy response genes. Onco Targets Ther 2018;11:7205–11.

35. Lu Y, Lin Q, Lin C, Chen J, Jiang X, He H. Down-regulation of miR-424 inhibited the metastasis of endometrial carcinoma via targeting PTEN/PI3K/AKT signaling pathway. J Cancer 2023;14(15):2811–19.

36. Mirantes C, Eritja N, Dosil MA, et al. An inducible knockout mouse to model the cell-autonomous role of PTEN in initiating endometrial, prostate and thyroid neoplasias. Dis Model Mech 2013;6(3):710–20.

37. Eritja N, Llobet D, Domingo M, et al. A novel three-dimensional culture system of polarized epithelial cells to study endometrial carcinogenesis. Am J Pathol 2010;176(6):2722–31.

38. Eritja N, Felip I, Dosil MA, et al. A Smad3-PTEN regulatory loop controls proliferation and apoptotic responses to TGF-beta in mouse endometrium. Cell Death Differ 2017;24(8):1443–58.

39. Forrest ARR, Kanamori-Katayama M, Tomaru Y, et al. Induction of microRNAs, mir-155, mir-222, mir-424 and mir-503, promotes monocytic differentiation through combinatorial regulation. Leukemia 2009;24(2):460–66.

40. Cancer Genome Atlas Research N, Kandoth C, Schultz N, et al. Integrated genomic characterization of endometrial carcinoma. Nature 2013;497(7447):67-73.

41. Rissland OS, Hong S-J, Bartel DP. MicroRNA Destabilization Enables Dynamic Regulation of the miR-16 Family in Response to Cell Cycle Changes. Molecular cell 2011;43(6):993–1004.

42. Eritja N, Navaridas R, Ruiz-Mitjana A, et al. Endometrial PTEN Deficiency Leads to SMAD2/3 Nuclear Translocation. Cancers (Basel) 2021;13(19).

43. Dosil MA, Mirantes C, Eritja N, et al. Palbociclib has antitumour effects on Pten-deficient endometrial neoplasias. J Pathol 2017;242(2):152–64.

44. Bartel DP. MicroRNAs: target recognition and regulatory functions. Cell 2009;136(2):215–33.

45. Subramanian A, Tamayo P, Mootha VK, et al. Gene set enrichment analysis: a knowledge-based approach for interpreting genome-wide expression profiles. Proc Natl Acad Sci U S A 2005;102(43):15545–50.

46. Peng Y, Croce CM. The role of MicroRNAs in human cancer. Signal Transduct Target Ther 2016;1:15004.

47. Braza-Boils A, Salloum-Asfar S, Mari-Alexandre J, et al. Peritoneal fluid modifies the microRNA expression profile in endometrial and endometriotic cells from women with endometriosis. Hum Reprod 2015;30(10):2292–302.

48. Haikalis ME, Wessels JM, Leyland NA, Agarwal SK, Foster WG. MicroRNA expression pattern differs depending on endometriosis lesion type. Biology of reproduction 2018;98(5):623–33.

49. Ohlsson Teague EM, Van der Hoek KH, Van der Hoek MB, et al. MicroRNA-regulated pathways associated with endometriosis. Mol Endocrinol 2009;23(2):265–75.

50. Rodriguez-Barrueco R, Nekritz EA, Bertucci F, et al. miR-424(322)/503 is a breast cancer tumor suppressor whose loss promotes resistance to chemotherapy. Genes Dev 2017;31(6):553–66.

51. Nekritz EA, Rodriguez-Barrueco R, Yan KK, et al. miR-424/503 modulates Wnt/beta-catenin signaling in the mammary epithelium by targeting LRP6. EMBO Rep 2021;22(12):e53201.

52. Rodriguez-Barrueco R, Latorre J, Devis-Jauregui L, et al. A microRNA Cluster Controls Fat Cell Differentiation and Adipose Tissue Expansion By Regulating SNCG. Adv Sci (Weinh) 2022;9(4):e2104759.

53. Langmead B, Salzberg SL. Fast gapped-read alignment with Bowtie 2. Nat Methods 2012;9(4):357–9.

54. Love MI, Huber W, Anders S. Moderated estimation of fold change and dispersion for RNA-seq data with DESeq2. Genome biology 2014;15(12):550.

55. Ewels P, Magnusson M, Lundin S, Kaller M. MultiQC: summarize analysis results for multiple tools and samples in a single report. Bioinformatics 2016;32(19):3047–8.

56. Patro R, Duggal G, Love MI, Irizarry RA, Kingsford C. Salmon provides fast and bias-aware quantification of transcript expression. Nat Methods 2017;14(4):417–19.

57. Soneson C, Love MI, Robinson MD. Differential analyses for RNA-seq: transcript-level estimates improve gene-level inferences. F1000Res 2015;4:1521.

58. Liberzon A, Subramanian A, Pinchback R, Thorvaldsdottir H, Tamayo P, Mesirov JP. Molecular signatures database (MSigDB) 3.0. Bioinformatics 2011;27(12):1739–40.

59. Walter W, Sanchez-Cabo F, Ricote M. GOplot: an R package for visually combining expression data with functional analysis. Bioinformatics 2015;31(17):2912–4.

60. Navaridas R, Vidal-Sabanes M, Ruiz-Mitjana A, et al. In Vivo Intra-Uterine Delivery of TAT-Fused Cre Recombinase and CRISPR/Cas9 Editing System in Mice Unveil Histopathology of Pten/p53-Deficient Endometrial Cancers. Adv Sci (Weinh) 2023;10(32):e2303134.

61. Eritja N, Mirantes C, Llobet D, et al. Long-term estradiol exposure is a direct mitogen for insulin/EGF-primed endometrial cells and drives PTEN loss-induced hyperplasic growth. Am J Pathol 2013;183(1):277–87.

