## Supplementary Tables S3-S7 for "Endometrial cancer progression driven by PTEN-deficiency requires miR-424(322)^∼^503"

**Table S3:** Genotyping primers and PCR protocols.

| **Allele** | **Primers** | | **PCR conditions** | | | **Product** |
| --- | --- | --- | --- | --- | --- | --- |
|  |  |  | **Temperature** | **Time (min)** | **Cycles** |  |
| **Cre:ER^T^** | Fwd: | ACGAACCTGGTCGAAATCGTGCG | 94 ºC | 2 min | 1 | Cre:ER^T-/-^: no band |
|  |  |  | 94 ºC | 45 sec | 32 |  |
|  |  |  | 65 ºC | 45 sec |  |  |
|  | Rev: | CGGTCGATGCAACGAGTGATGAG | 72 ºC | 45 sec |  | Cre:ER^T+/-^: 350 bp |
|  |  |  | 72 ºC | 5 min | 1 |  |
|  |  |  | 4 ºC | ∞ | |  |
| ***Pten*^f/f^** | Fwd: | CAAGCACTCTGCGAACTGAG | 94 ºC | 3 min | 1 | *Pten*^+/+^: 156 bp |
|  |  |  | 94 ºC | 30 sec | 35 |  |
|  |  |  | 60 ºC | 1 min |  | *Pten*^f/+^: 156 and 328 bp |
|  | Rev: | AAGTTTTTGAAGGCAAGATGC | 72 ºC | 2 min |  |  |
|  |  |  | 72 ºC | 2 min | 1 | *Pten*^f/f^*: 328* bp |
|  |  |  | 4 ºC | ∞ | |  |
| ***Smad2*^f/f^** | Fwd: | TGAGACTTCTCTGTACCCGAT | 95 ºC | 5 min | 1 | *Smad2*^+/+^: 350 bp |
|  |  |  | 95 ºC | 30 sec | 40 |  |
|  |  |  | 58 ºC | 45 sec |  | *Smad2*^f/+^: 350 and 400 bp |
|  | Rev: | CATCAGATTCCATTAGAGATGG | 72 ºC | 45 sec |  |  |
|  |  |  | 72 ºC | 7 min | 1 | *Smad2*^f/f^: 400 bp |
|  |  |  | 4 ºC | ∞ | |  |
| ***Smad3*^f/f^** | Fwd: | CTCCAGATCGTGGGCATACAGC | 95 ºC | 2 min | 1 | *Smad3*^+/+^: 150 bp |
|  |  |  | 95 ºC | 30 sec | 30 |  |
|  |  |  | 60 ºC | 45 sec |  | *Smad3*^f/+^: 150 and 200 bp |
|  | Rev: | GGTCACAGGGTCCTCTGTGCC | 72 ºC | 45 sec |  |  |
|  |  |  | 72 ºC | 10 min | 1 | *Smad3*^f/f^: 200 bp |
|  |  |  | 4 ºC | ∞ | |  |
| **miR-322/503^+/+^** | Fwd: | CACCAGCAGATCCTGGAAAT | 94 ºC | 2 min | 1 | miR-322/503^+/+^: 500 bp |
|  |  |  | 94 ºC | 30 sec | 35 |  |
|  |  |  | 59 ºC | 30 sec |  |  |
|  | Rev: | CAAGTGAGGCGCTAACAACA | 72 ºC | 2 min |  | miR-322/503^-/-^: no band |
|  |  |  | 72 ºC | 5 min | 1 |  |
|  |  |  | 4 ºC | ∞ | |  |
| **miR-322/503^-/-^** | Fwd: | AGTTTCGAAAAACGGACATA | 94 ºC | 2 min | 1 | miR-322/503^+/+^: no band |
|  |  |  | 94 ºC | 30 sec | 35 |  |
|  |  |  | 59 ºC | 30 sec |  |  |
|  | Rev: | ATTGCCTCTCATTGTACCAC | 72 ºC | 2 min |  | miR-322/503^-/-^: 700 bp |
|  |  |  | 72 ºC | 5 min | 1 |  |
|  |  |  | 4 ºC | ∞ | |  |

**Table S4:** Primers and conditions used for RT-qPCR.

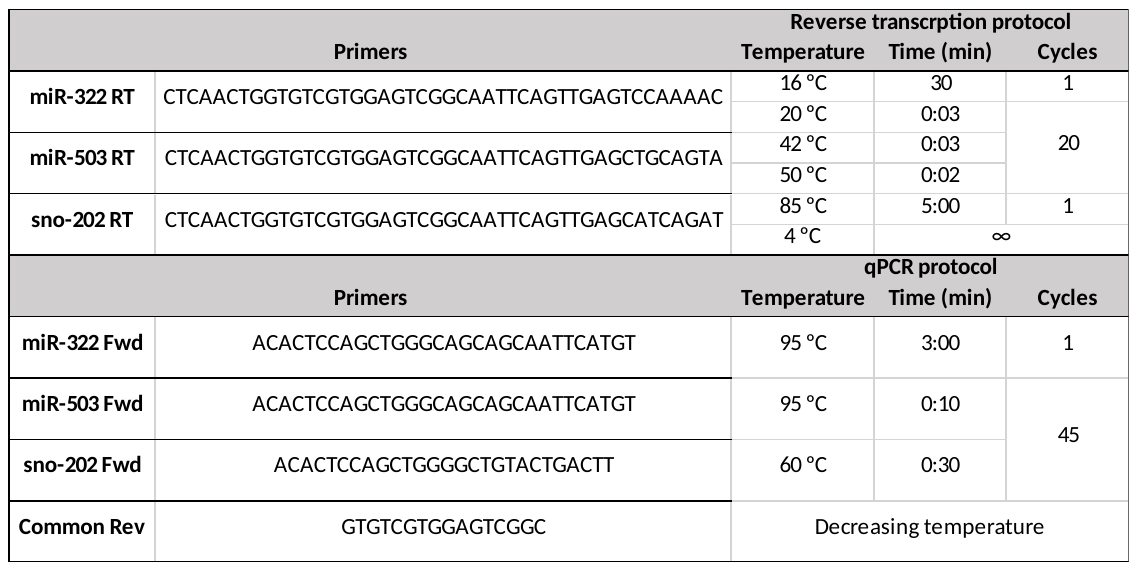

**Table S5:** Antibodies and dilutions used for Western Blot.

| Antigen | Dilution | Catalogue number, Company |
| --- | --- | --- |
| GAPDH | 1:10000 | 8245, Abcam |
| Cyclin D1 | 1:1000 | SC20044, Santa Cruz |
| p-AKT (Ser473) | 1:1000 | 4060, Cell Signaling |
| p-p70SK (Thr389) | 1:1000 | 9205, Cell Signaling |
| p-S6 (Ser235/236( | 1:1000 | 2211, Cell Signaling |
| p-Tsc2 (Thr389) | 1:1000 | 3611, Cell Signaling |
| panERK | 1:1000 | 610623, BD Biosciences |
| p-IGF1Rβ (Tyr1135/1136) | 1:1000 | 3024, Cell Signaling |
| PTEN | 1:1000 | 9188, Cell Signaling |
| Smad3 | 1:1000 | SC101154, Santa Cruz |
| Mouse IgG HRP | 1:10000 | 115-135-003, Jackson |
| Rabbit IgG HRP | 1:10000 | 111-035-003, Jackson |
| Goat IgG HRP | 1:10000 | 705-035-147, Jackson |

**Table S6:** Antibodies and dilutions used for BrdU and immunofluorescence.

| Antigen | Dilution | Catalogue number, Company |
| --- | --- | --- |
| BrdU | 1:100 | 56258, Santa Cruz Biotechnology |
| Cleaved caspase 3 (Asp175) | 1:250 | 9661, Cell Signaling |
| Phalloidin | 1:500 | P5282, Sigma |
| Mouse IgG Green (Alexa Fluor™ 488) | 1:500 | A11029, ThermoFisher |
| Rabbit IgG Green (Alexa Fluor™ 488) | 1:500 | A11008, ThermoFisher |
| Rat IgG Green (Alexa Fluor™ 488) | 1:500 | A11006, ThermoFisher |

**Table S7:** Antibodies and dilutions used for immunohistochemistry.

| Antibody | Dilution | Catalogue number, Company | Antigen Retrieval | Secondary Antibody |
| --- | --- | --- | --- | --- |
| PTEN | 1:100 | M3627, DAKO | High pH | EV Flex kit |
| p-AKT (Ser473) | 1:50 | 3787, Cell Signaling | High pH | Anti-rabbit biotin |
| Anti-mouse biotin | 1:200 | 115-065-166, Jackson | - | - |
| Anti-rabbit biotin | 1:200 | 111-065-144, Jackson | - | - |
| Estreptavidin HRP | 1:400 | P0397, Dako | - | - |
